# Focus on the edges: a biomolecular network of histone PTMs, metabolites and proteins unveils functional entanglement in AML

**DOI:** 10.64898/2026.04.22.716296

**Authors:** Boris Vandemoortele, Laura Corveleyn, Jolien Vanhooren, Amélie De Maesschalck, Jolien De Waele, Thijs Lefever, Ruben Almey, Ivo Kwee, Antonino Zito, Larissa Deneweth, Simon Daled, Barbara De Moerloose, Dieter Deforce, Vanessa Vermeirssen, Tim Lammens, Maarten Dhaenens

## Abstract

The cell phenotype is not a direct manifestation of the genotype but rather a product of cellular history and the environmental context. However, individual biomolecules cannot change independently and show coordinated behavior. To study this in acute myeloid leukemia (AML), we built a unique multi-omics biomolecular network made from proteins, metabolites and histone posttranslational modifications (hPTMs) sequentially extracted from each cell pellet. Edges between the nodes are measured directly using 400 LC-MSMS runs that cover 18 AML cell lines. We provide a novel conceptual framework to illustrate the different classes of functional entanglement between and within omics layers and present the data in three interactive data browsers to allow full community access. To help navigate the network, we approach it from the perspective of two biomolecular targets, i.e. CD34 and the epigenetic mark Histone H3 lysine 27 trimethylation (H3K27me3). Now, this easily accessible biomolecular network serves as a starting point for building and testing hypotheses and streamlining drug development, in the process positioning biomolecular associations center stage in understanding phenotypic complexity.

## INTRODUCTION

Attributing traits or diseases solely to genetic aberrations has proven to be an oversimplification ^1,2^. Even protein function alters drastically as a product of its highly dynamic environment, i.e. subcellular localization, cellular context, and surrounding tissue, and therefore cannot be solely encoded in the genomic sequence. Additionally, the complex interactions with neighboring proteins and metabolites together lead to higher-level molecular networks, i.e. macroscales, that boost biological resilience, most notably in Eukaryotes ^3^. It is therefore hypothesized that multicellular complexity was attained not through a proportional increase in the number of protein-coding genes (i.e. nodes in the network), but rather by the exploitation of the combinatorial space, i.e. increasing the number of biomolecular interactions. Therefore, while the nucleotide-centric perspective and the loss-of-function phenotype approaches commonly used in molecular biology remain foundational, they have not been able to fully capture how phenotypes emerge in health and disease.

Interestingly, purely non-genetic, epigenetic mechanisms were recently found to be sufficient to initiate tumorigenesis in stem cells, irrespective of genomic aberrations. *Parreno et al.* showed that a transient loss of transcriptional silencing by Polycomb group proteins (PRC2) that mediate histone H3 lysine 27 trimethylation (H3K27me3) was sufficient to induce an irreversible switch to an epigenetic cancer cell fate^3^. The same hPTM in fact is a gatekeeper of stemness in both human and mouse embryonic stem cell biology ^4–6^. Interestingly, stemness is a driver mechanism in many cancers ^7^. Acute Myeloid Leukemia (AML) for example, is a heterogeneous hematologic disease that is characterized by unrestricted proliferation of abnormal myeloid precursor cells within the bone marrow, which results in impaired hematopoiesis. While genomic studies have established driver events in AML, some of which constitute prognostic subgroups, a recent study by Jayavelu et al. identified a novel clinical subtype of AML (Mito-AML) which could only be detected through proteomic analysis, i.e. a phenotypic manifestation^8^. Mito AML is associated with higher mitochondrial activity and increased resistance to Venetoclax, a Bcl-2 inhibitor ^8^. The impact of these beyond-the-genome discoveries is further substantiated by the recent success of Menin (MEN1) inhibitors, such as Revuminib, targeting the histone epigenome to trigger a massive differentiation event reflected by a brief spike in cell counts (differentiation syndrome), after which leukemic cells die ^9^.

MS-based omics study functional biomolecules in a cell that together define the phenotype. The recent advances in MS instrumentation and analysis tools, combined with easily accessible databases that capture past experimental work and functional connections, now allow asking much broader conceptual questions beyond the binary comparison of cellular phenotypes, thus contributing to a deeper understanding of how life works. This becomes even more powerful when multiple phenotypic biomolecular classes are integrated through a multi-omics approach and when this is applied to more complex experimental designs. This allows to build robust biomolecular networks, with the different biomolecules as the nodes and their correlations creating the edges. Overall, such networks shift the focus from single gene products to biological modules, which are now integrated as fundamental features of the data architecture, extending on the concept of “macroscales” defined by protein-protein interactions ^3^. Indeed, by building the network with MS-based omics, objective and robust, yet more distant correlations between abundances emerge. We therefore propose to extent this concept to “functional entanglement”, now including coherent biomolecular changes that appear to have no first order causal connection, as these underly Eukaryote complexity. This is the directly measured, multi-omics equivalent of more overarching and public data efforts described elsewhere ^10^.

Here, we develop a novel multi-omics approach that enables detecting and studying functional entanglement. The power of the approach lies in the unique combination of the three functionally entangled omics layers, i.e. the histone epigenome, the metabolome and the proteome that are sequentially isolated from the same samples ^11–13^. The complex experimental design in total comprises more than 400 LC-MSMS runs, i.e. three omics layers of six replicates of 18 different unputurbed AML cell lines. This data matrix allows quantifying robust associations within and between omics layers to find co-regulated biomolecules through the interplay between metabolic state, epigenetic read-out and the expressed protein pool. As this conceptually can be studied for every biomolecule in the network, we necessarily provide the data in a community-accessible way for others to investigate functional entanglement from the perspective of their biomolecular target of interest to develop follow-up mechanistic studies: (i) the biomolecular network built through Lemonite ^14^, (ii) the uniquely inferred hPTM abundances statistically processed through differential usage ^15,16^ and (iii) the proteome and multi-omics datasets accessible as over 120 interactive plots, that can be created for all biomolecules in a Omics Playground (Bigomics) project. The dozens to hundreds of existing studies that already describe many of the edges captured in the network, provide an estimate of the prior probability in selecting the novel, unknown connections for mechanistic follow-up in the future.

We facilitate navigation of this intricate multi-modal network by providing a conceptual biological framework in the form of a circular graphical path diagram that illustrates the information content of each of the layers (**Figure 1A**). First, we describe the overarching biomolecular network and explain how it can be functionally interpreted. As an illustration, we interrogate a tiny corner of the data space, i.e. the most densely connected node and stemness marker Histone H3 lysine 27 trimethylation (H3K27me3) and its known anti-correlating Histone H4 lysine 8 acetylation (H4K8ac), while leveraging the prominent AML biomarker CD34 to illustrate the network’s coherence. Subsequently, we introduce functional entanglement and illustrate how the highest scoring edge weights often provide evidence of protein complexes, inferring interactomics data without a dedicated protocol. As this is induced by selective degradation of unbound protein units, we propose the term protein complex dosage compensation, as the proteomics equivalent of the gene term coined elsewhere ^17^. Next, we explore the underlying omics layers separately to illustrate the value of the edges in each of the layers, and we end by demonstrating how inter-omics correlations allow investigating overarching molecular behaviour, extending functional entanglement to include both first and n^th^ order causal interactions. In other words, we define the edges rather than the nodes as the most fundamental building blocks of Eukaryote complexity and the resulting cellular phenotypes, because cancer continuously explores “new” cellular phenotypes from the same genotype that no longer obey the checks and balances of normal development and overgrow the healthy organism. This is the first comprehensive knowledge base that captures functional entanglements of all orders over these three biomolecular classes and will help streamline drug development in the future.

**Figure 1.**
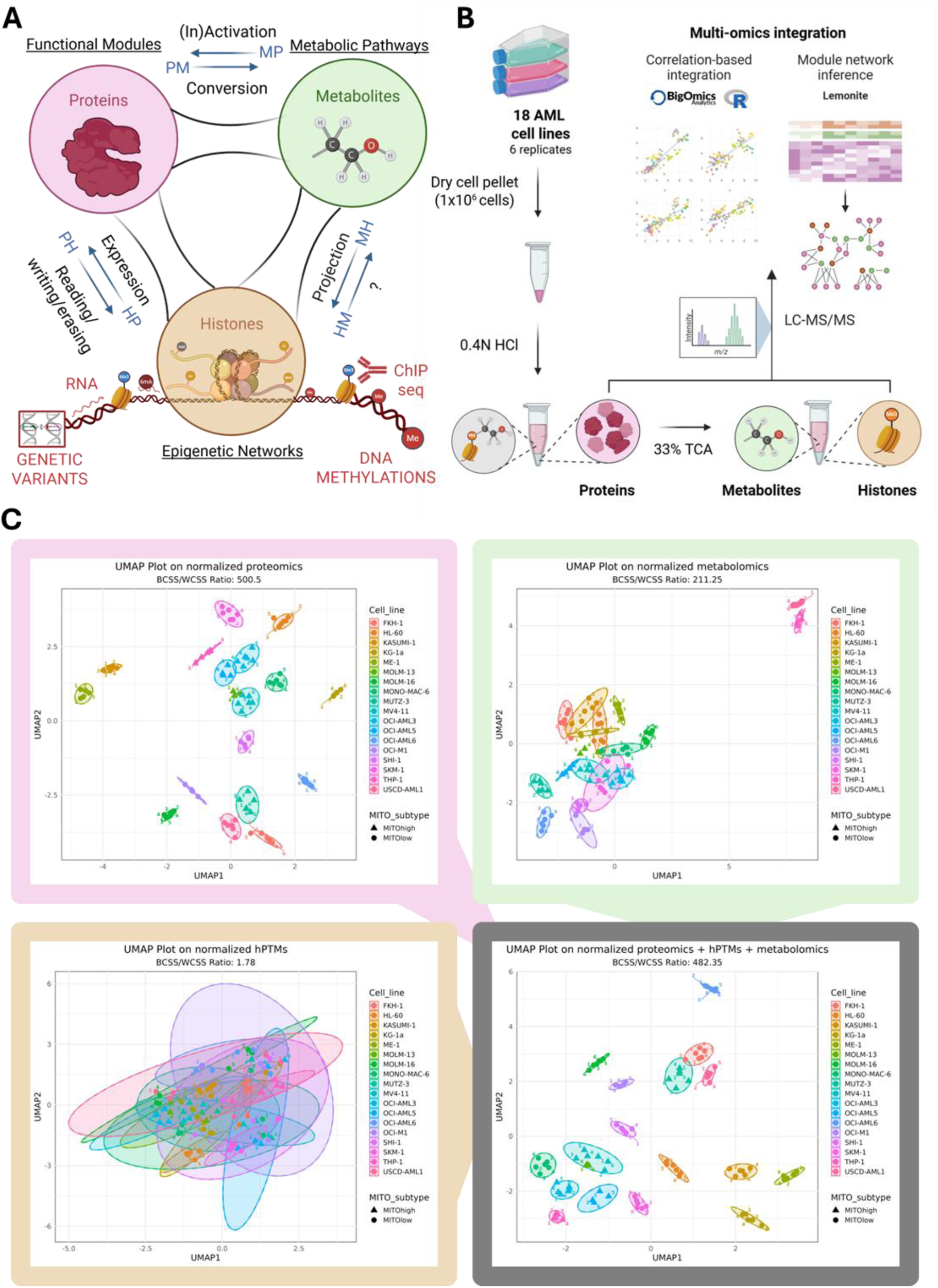
Conceptual framework. **A. The Biology perspective**. Cancer biology has been mainly studied through integration of sequencing technologies, amongst others targeting DNA mutations/variants, DNA methylations, histone PTMs by ChIPseq, and (non-) protein-coding RNA expression (bottom, in red). However, the phenotype is a product of selective activation of different parts of the genome throughout cell and tissue history. The conceptual framework in our study centres on a perspective of interactions between histones, proteins and metabolites that defines the phenotype. Co-regulated functional protein modules, metabolic pathways and epigenetic networks integrate to define function. Starting from an external stimulus, histones will help modify chromatin structure and alter protein expression (**HP**). Some of these proteins like readers, writers and erasers will directly modify hPTMs (**PH**) and others will convert metabolites to harvest available energy sources (**PM**). Some of these metabolites can modify protein activity, e.g. as co-factors) (**MP**) and others can directly modify hPTMs or activate/inhibit their readers, writers and erasers (**MH**). This view implies that inversely, hPTMs could become metabolites, which we confirmed in our data. **B.** The Protocol Perspective. We developed a minimalistic workflow for sequential extraction of the proteome, histonome and metabolome and performed it on 6 biological replicates of 18 different AML cell lines. These three distinct biological fractions were processed individually for LC-MS/MS analysis and measured in three fully randomized quality-controlled sample batches. HCl: Hydrochloric Acid; TCA Trichloric acid. **C.** UMAPs of all the identified features in each of the omics layers (color-coded according to **A**). Grey-lined UMAP is integrated from all biomolecular features in a single multi-omics UMAP. All cell lines can be completely separated, illustrating apt analytical space for studying edges, which are built from co-ordinated behaviour of biomolecules within and between omics layers.

## RESULTS

### The conceptual, biological and experimental framework

The power of multi-omics analyses has been illustrated over the past decade, mainly through integration of sequencing technologies, amongst others assessing DNA variations, DNA methylation, histone PTMs by ChIPseq/Cut&TAG, (non-)-coding RNA expression and posttranscriptional modifications (**Figure 1A**, red at the bottom). However, other omics layers further manifest the biomolecular phenotype, including, but not limited to, the assembled proteome, metabolome and histone posttranslational modification (hPTM) repertoire. While these have separately been studied by mass spectrometry (MS) in cancer biology, they have never been measured before in a single experiment, let alone from a single sample. Yet, doing so uniquely allows quantifying their interactions, i.e. the edges in the biomolecular network.

First, we propose a novel circular graphical path diagram that depicts the potential functional relationships in molecular biology that could be probed within this dataset (**figure 1A**). Briefly, cellular and environmental cues are transferred to the nucleus, causing the activation or inactivation of specific genomic regions by altering the hPTM repertoire. This changes RNA transcription (**Figure 1A_HP**), altering the available protein repertoire. Some of these proteins can e.g. mediate epigenetic changes (writers and erasers) and interpret them (readers) to further alter protein expression (**Figure 1A_PH**), while others convert metabolites to harvest energy and create new building blocks (**Figure 1A_PM**). The latter process not only can generate energy but can equally change the molecular context of the proteins, thus changing their function or simply inactivating or activating them by metabolite co-factors (**Figure 1A_MP**). More recently, it became accepted that these metabolites can equally get projected onto histones (non-) enzymatically^11^ (**Figure 1A_MH**) ^11^, where they effectively translate the energy state of the cell into a molecular fingerprint that defines the protein expression repertoire through epigenetics (**Figure 1A_HP**). We refer to the latter edge as the “Energy-Information Axis”, closing a circular interaction between histones, proteins and metabolites that captures many of the changes that define the phenotype. At the onset of this study, the existence of metabolites derived from histones was unknown to us, yet our data pinpoints towards at least two such metabolites, i.e. trimethyllysine and dimethylarginine (**Figure 1A_HM**). **Supplementary Figure 1** illustrates how the interactions between the three biomolecular layers are currently interpreted manually, here from the perspective of one of the most densely connected hPTMs in the biomolecular network and a known stemness epigenetic mark, i.e. trimethylation of lysine 27 of histone H3: **H3K27me3**.

**Figure 1B** displays the minimalistic protocol that was developed to measure the proteome, hisonte PTMs and metabolome from a single sample. To generate a proof-of-principle unperturbed dataset, we collected cell pellets (2.000.000 cells) of 18 genetically different AML cell lines (6 replicates each). From the same cell pellet, samples were extracted and prepared for metabolomics, hPTM analysis and proteomics. All samples were measured in four independent, fully randomized and quality-controlled sample batches (metabolomics were measured once on C18 and once on HILIC and depicted as _C18 and _HI, respectively). This creates an extremely coherent and low-noise data matrix. Only samples without outliers in any of the omics layers were retained, resulting in a final dataset of 97 samples. Encompassing over 150 hPTMs (from ∼500 peptidoforms), 4500 reproducibly detected proteins (from about 70.000 peptides), and over 700 confident metabolites (out of 15.000 features), this dataset provides an exceptionally comprehensive image of the molecular phenotype of 18 different AML genotypes. QC reports are available in **Supplementary Data 1**.

This dataset provides a snapshot at one point in the cellular history of hitherto undisturbed cells, further annotated based on extensive available metadata (**Supplementary Data 1**). To quantify the edges in the network requires numerical data, which here is provided by the measured abundances of the biomolecules and their differences between the different genotypes. Exploratory UMAPs comprehensively capture the differences per omics layer and across modalities (Figure 1C). The quality control (QC) samples are centrally located in all the UMAP plots, confirming instrument stability throughout the three biomolecular batches measured in this study. Replicates of cell lines cluster more tightly together based on proteome and metabolome data compared to hPTM data. To date, very few studies have applied MS to measure overall hPTM abundance in such complex experimental designs, and we cannot therefore verify if this is a consistent finding. Still, we hypothesize that this inherent lower separating power of hPTM data derives from (i) the smaller fold changes that occur genome-wide, (ii) the lower number of final features in this data layer and (iii) the higher complexity of sample preparation and data analysis for hPTM analyses ^15,18–21^.

Three separate interactive reports are provided for easy interrogation of this extensively preprocessed data at http://www.lemonite.ugent.be/AML_Results:

1. The comprehensive interactive multi-omics module network created through Lemonite ^14^, where protein modules are connected to potential “regulatory” hPTMs and metabolites, genetic and clinical metadata, and functional enrichment analysis.
2. The interactive html histone analysis report, documenting normalization strategies, histone variant inferences, peptidoforms and hPTM differential usage ^15^, extendable to extract custom contrasts.
3. The pre-processed proteomics and multi-omics data provided as Omics Playground (Bigomics) projects, comprising over interactive 120 plots where every protein target can be investigated separately. Register at Bigomics and find these under public datasets.

The full set of available files and analyses is summarized in a MindMap under **Data availability**. This dataset comprises 5521x5520 quantified edges, i.e. over 30 million edges from 5521 nodes. This is the theoretical equivalent of how Eukaryotes leveraged interactions to attain complexity. These can all be investigated through three interactive webbrowsers/frameworks by non-bioinformatics experts, making it the most easily accessible and interpretable multi-omics dataset to date that directly captures the biomolecular phenotype of this many cancer cell lines. Therefore, a video tutorial with brief biological introduction (until minute 15) is available online: https://youtu.be/pSfDwsrRzm0. As a guide to navigate this data, we investigate in depth the multi-omics perspective from a single biomolecular target that has not been extensively studied in AML, i.e. **H3K27me3**.

### A data-driven and interpretable biomolecular network on 18 AML cell lines

In search of biomarkers for a given disease or treatment, omics studies have long focused on separate abundances of biomolecules, often in experimental designs of binary comparisons. Because life seems to operate at higher levels, however ^1,3,22^, others and we advocate for a shift towards studying associations between biomolecule abundances, rather than focussing on the biomolecules themselves. With an adequately complex experimental design, such interactions can be readily calculated as robust correlations, which in turn can be visualized as edges in a network where the nodes are the biomolecules. In other words, we here study the edges of the biomolecular network, rather than the nodes.

#### The biomolecular network unveils functional entanglement

Lemonite, a probabilistic multi-omics module network inference framework that combines ensemble clustering of biomolecules into modules with an ensemble of decision trees to assign potential “regulators” to each module ^14^. Herein, “regulators” is used as a data term, and this does not necessarily imply biological causality or directionality. This method provides a single data-driven network view of the biomolecular phenotype and is the full integration of the conceptual framework in Figure 1A. We chose the richest annotated omics layer, i.e. the proteins, to create the initial modules for the network. Modules for the hPTMs and metabolites were equally created and can be found separately in **Supplementary Data 2.** The network is built up from the proteomics layer where 2521 proteins (out of 4612 proteins) were grouped into 68 protein modules (blue circles), with x metabolites (yellow) and y hPTMs (blue) attributed to them as “regulators” with specific scores reflecting the relevance of their association to the protein module (Figure 3A). A visually attractive, interpretable and interactive web browser displays this multi-omics network at http://www.lemonite.ugent.be/AML_Results. **Supplementary Table x** provides a summary of the network.

**Table 1.** Overview of Mascot search parameters.

|  | Quality search | Error tolerant search | Biological search |
| --- | --- | --- | --- |
| Mass error precursor ions (ppm) | 10 | 10 | 10 |
| Mass error fragment ions (ppm) | 50 | 50 | 50 |
| Enzyme specificity | Trypsin | ArgC | ArgC |
| Miscleavages | 1 | 1 | 1 |
| Variable modifications | Propionylation (N-term)<br>Propionylation (K) | Deamidation (NQ)<br>Oxidation (M) | Acetylation (K)<br>Butyrylation (K)<br>Lactylation (K)<br>Trimethylation (K)<br>Succinylation (K)<br>Methylation (R)<br>Dimethylation (KR)<br>Deamidation (NQR)<br>Phosphorylation (ST) |
| Fixed modifications | / | Propionylation (N-term)<br>Propionylation (K) | Propionylation (N-term)<br>Propionylation (K) |
| Database | Human Swissprot supplemented with contaminants from the common Repository for Adventitious Proteins (cRAP) database | Human Swissprot supplemented with contaminants from the common Repository for Adventitious Proteins (cRAP) database | Custom-made FASTA-database |

The edges quantify functional relationships and thus provide the broader biomolecular context in which each biomolecule is operating. At the protein level, co-expressed proteins are grouped in modules and their functional coherence can be assessed by gene set enrichment analysis (GSEA). In the Lemonite network, all proteins in the network are ranked based on Pearson correlation to module eigengene expression. Thus, whenever a given regulator is higher in abundance, a given protein module is up- or downregulated too, in turn implying that its function is up- or downregulated in the cellular context of the respective regulator, without implying any causality *per se*. Rather, the regulator cannot be seen uncoupled from this function and its connection to the protein module function therefore reflects a “**functional entanglement**”. For the hPTM and metabolite regulators these functional entanglements will provide a crucial knowledge base during drug development.

#### The biomolecular network topology

The network contains 355 nodes and 740 edges. Figure 3A depicts the network, highlighting several of the highest regulator-module edge weights shown in Table 1, plus the rest of the interconnected modules to these regulators. Topological analysis reveals that the regulator outdegree follows a scale-free distribution (γ = 1.74 and R² = 0.935), suggesting the network’s behaviour is fundamentally driven by a set of central regulatory hubs. Notably, the same analysis on the expanded network with all proteins as separate nodes showed that the regulator outdegree is random (γ = 0.5), illustrating how the prior clustering into modules exposes the true regulatory skeleton.

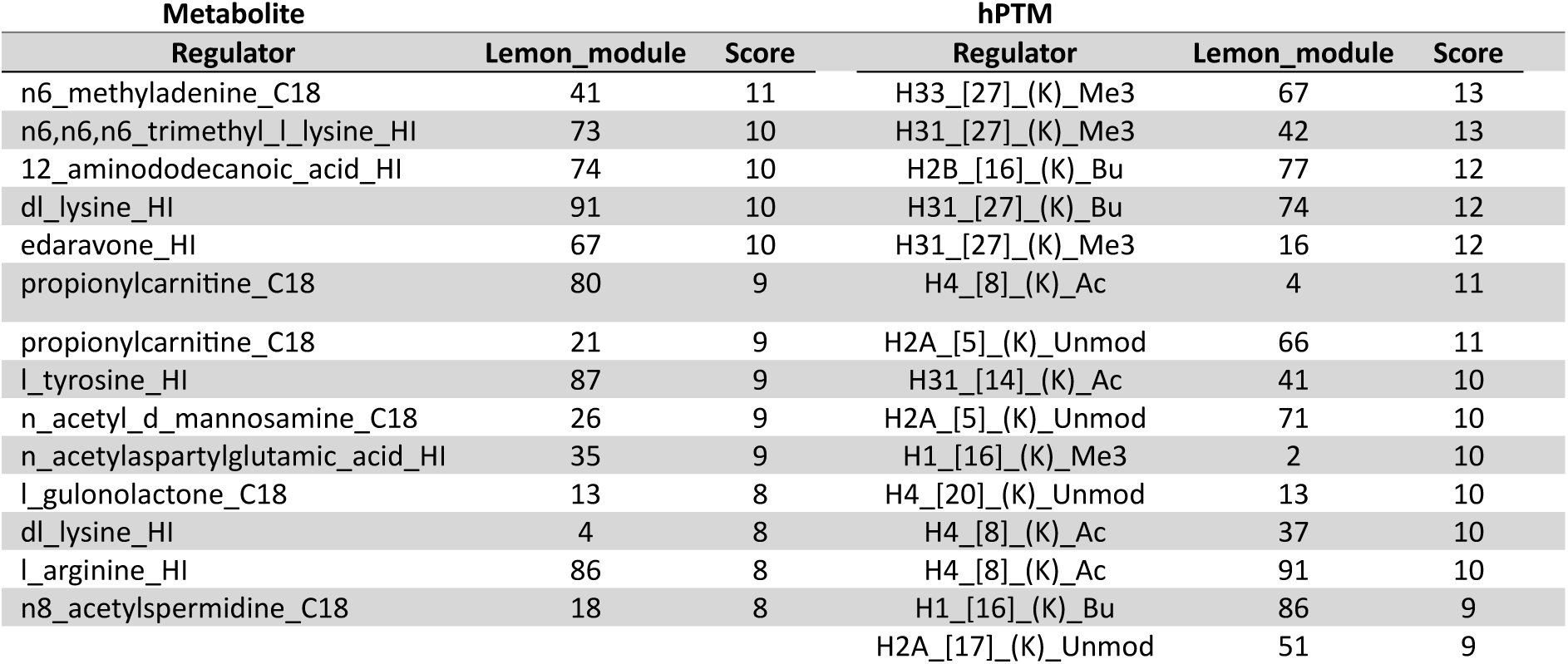

For histones, **H3K27** makes up the top four strongest edges, with trimethylation (H3K27me3) linking to module 67, 42 and 16, and monomethylation (detected as butyrylation in our workflow) linking to module 74. **Figure 3A and 3B** depict module 16, and Figure 3C shows the GSEA results from Lemonite, which is enriched for *structural constituent of the nuclear pore.* Members of this protein complex that are enlisted in the CORUM protein complex database are highlighted in orange. These correlations can then be investigated in detail using the Omics Playground (Bigomics) project. Figure 3D shows the five highest correlating proteins (out of all 4566) to nucleoporin 93 (NUP93) plotted over the 18 AML cell lines. Here too, these are other nuclear pore members as well as dynamin 2 (DNM2), which is crucial for vesicle trafficking from the trans-Golgi network. TRIM28, RCC1 and CTNNBL1 are ranked next in this list and are equally depicted in Figure 2. From this, and from the yellow band to the right of module 16, it is also clear that some correlations are inflated by a single cell line (MOLM-16 in this case), but that this captures biology, nonetheless. Another hPTM, acetylation of H4 lysine 8 (**H4K8ac**), is found three times in the histone regulator list top 11 (**Table 1**), connecting to module 4, 37 and 91 (Figure 3A). All three top-scoring modules are themselves functionally negatively associated to *Respiratory chain complex I* and positively associated to *vesicle organization* and *Golgi membrane*, with an additional positive correlation between module 37 and *MHC protein complex* (Figure 2). Remarkably, BRD4, an important reader of H4K8ac, also resides in module 37. It is therefore likely that this hPTM and its reader are stabilized by mutual binding, protecting each other from removal/degradation. This interpretation is the multi-omics equivalent of “protein complex dosage compensation”, whereby unincorporated subunits from a complex are rapidly degraded by the proteasome and protein complexes become strongly enriched in the modules. Module 37 also strongly negatively correlated with three metabolite regulators containing glutamine. In total, H4K8ac links to 12 modules. Further down the ranking, H4K8Ac is negatively correlated to module 72, which contains **mTOR**, a central regulator of cellular metabolism in response to nutrients, energy and stress signals. mTOR has been hypothesized to be the direct energy sensor for H3 and H4 acetylation for opening chromatin in times of abundant energy-availability ^11^.

**Figure 2.**
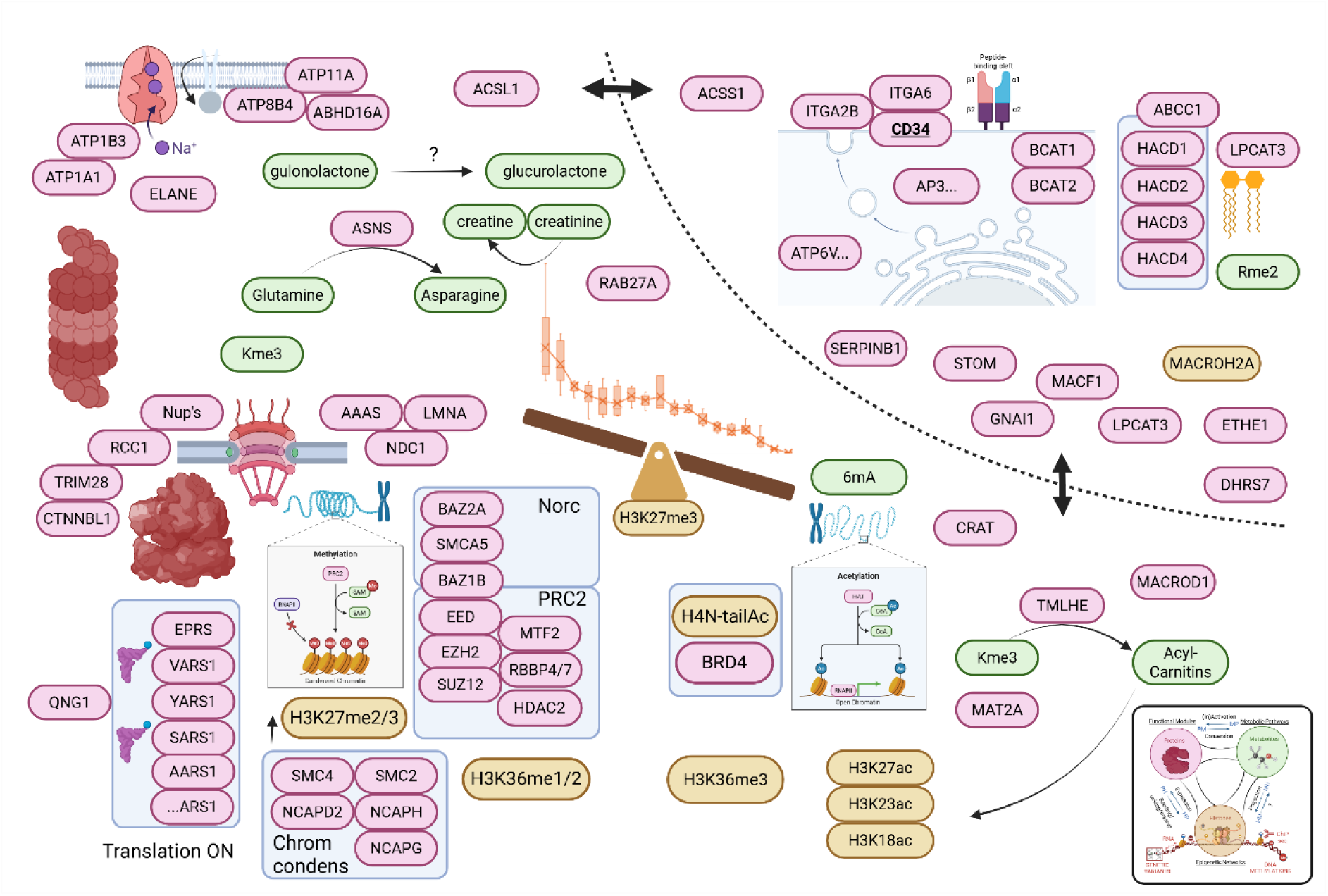
Biomolecular associations from the perspective of H3K27me3: an overview of the results. Correlations between and within the three biomolecular layers (proteins, metabolites and hPTMs color-coded according to the inset) are mined throughout the results section from the perspective of H3K27me3, guided by the conceptual framework depicted in **Figure 1A**. H3K27me3 abundances over the 18 AML cell lines from manually isolated and targeted example provided in **Supplementary Figure 1A** are centrally depicted from high to low and iconized into a balance to illustrate that all downstream analyses are not merely based on a binary contrast, but rather on correlations, i.e. the edges, that represent the protein, hPTM and metabolite abundances that consistently co- or anti-correlate with this hPTM. Those proteins that form known protein complexes are grouped in blue squares. In turn, these provide a glimpse of the potential phenotypes that can(not) exist in AML, because this data demonstrates how single biomolecules cannot independently change their abundance without all other biomolecules changing with it: when a biomolecule to the left increases, all other biomolecules to the left also increase and those to the right decrease, irrespective of causality. In other words, the changes depicted here are “unbreakable” within the AML cell line phenotype and are therefore functionally entangled. Note that this two-dimensional representation cannot capture all mutual (anti-)correlations between the molecules. As an example, the dotted line represents a more binary-like representation of the CD34-co-enriched (and downregulated) proteins from **Figure 4**.

From the metabolite perspective, **6-methyladenine (m6A)** is the most tightly connected metabolite, negatively correlating to module 41, which contains CD33 and prelamin A/C (LMNA). Dysregulation of the m6A machinery plays a crucial role in the initiation, progression, and maintenance of AML, particularly in the self-renewal of leukemic stem cells (LSCs)^23^. Recently, the RNA *N*^6^-methyladenosine (m^6^A) modification was found as a key regulator of serine biosynthesis in AML ^24^ and indeed three out of its modules (85, 41 en 24) are strongly enriched for selenoamino acid metabolism. Yet the role of the m6A metabolite itself is currently unknown. m6A derives from degradation of RNA containing methylated adenine residues, which is a postsynthetic modification by the S-adenosyl-L-methionine (SAM) methyl donor. It therefore could be considered a metabolic echo of a posttranscriptional modification. Next to **lysine**, which links to **H4K8ac** through module 4 (Figure 3A), **n6,n6,n6 trimethyllysine (Kme3)** also is amongst the top-ranking edges, tightly anti-correlating to module 73. Kme3 is released from histones and other proteins via proteolysis where it was equally generated by the action of SAM to form the trimethylation PTM (HMDB0001325 description). Once released in muscles, it serves as the precursor for carnitine biosynthesis and it is a coenzyme of fatty acid oxidation. Note that **Kme3** was also the highest correlating metabolite edge to module 16, which links most strongly to **H3K27me3** (**Figure 3A and 3B**). While never studied in leukaemia, this observation could be interpreted as a potential case of **HM directionality in** Figure 1A, i.e. a histone PTM that becomes a metabolite with downstream metabolic impact. Clearly, this potential metabolic echo of an epigenetic modification cannot be traced to H3K27me3 alone and most probably derives from a mixture of many other modified residues. Yet it is encouraging to also see two detections of **propionylcarnitine** in the top 10 most strongly connected metabolites, as regulator of module 21 and 80 (Figure 3A). Module 21 contains **MAT2A**, S-adenosylmethionine synthase isoform type-2, the enzyme that converts methionine into SAM. Additionally, **propionylcarnitine** inversely correlates to three modules of **H3K27me3**, i.e. 96, 66 and 81, while it correlates in the same sense as **H4K8ac** to modules 46 and 90, the latter anticorrelating with m6A. This closes the loop with a potential route feeding the protein/histone trimethylation, which then becomes Kme3 and later carnitine, as studied further below.

**Figure 3.**
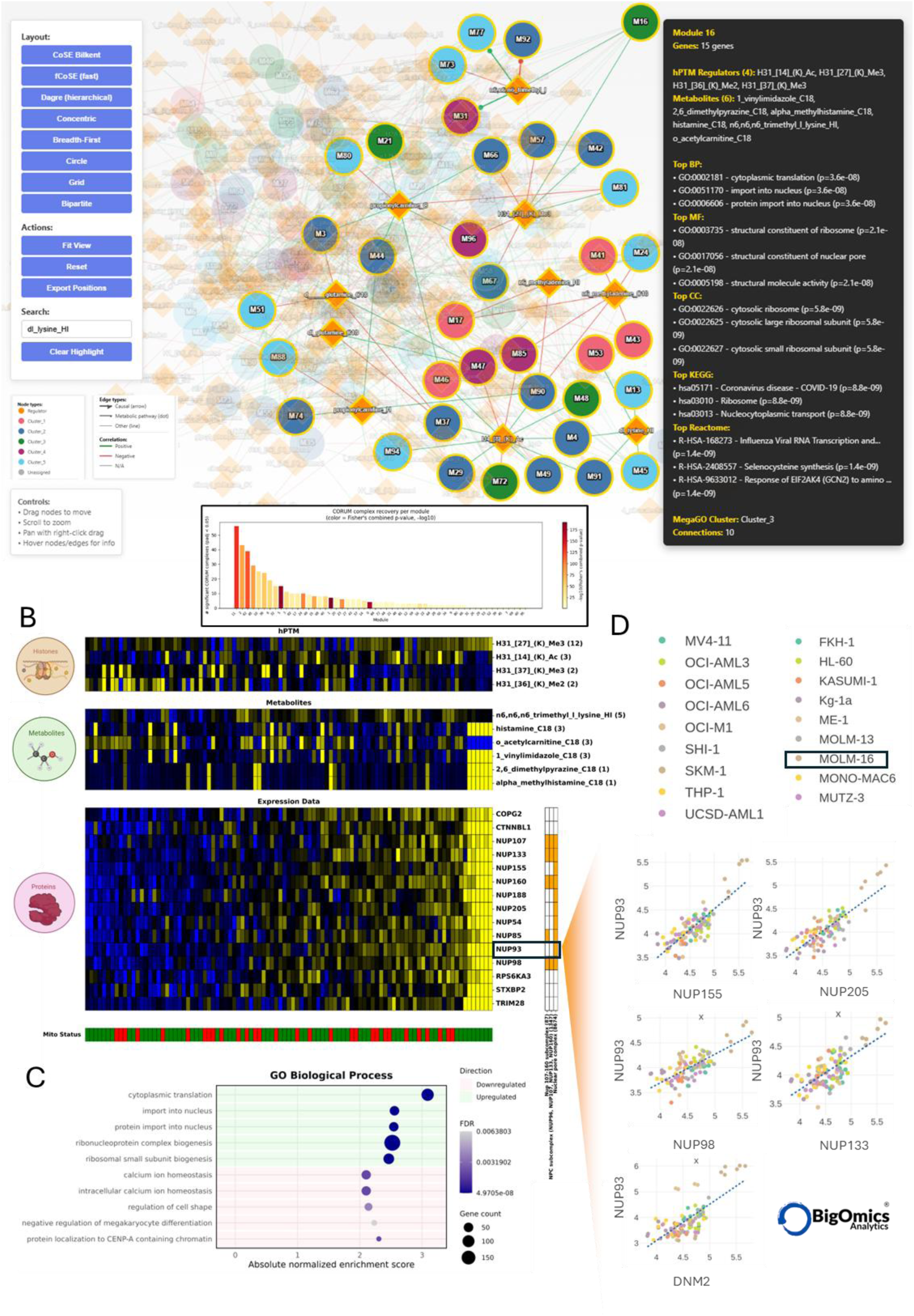
The Lemonite biomolecular network interactively accessible online. **A.** The protein data layer was used to create 68 modules which are connected to regulators, i.e. hPTMs and metabolites through an ensemble of decision trees. This provides a single, comprehensive readout of the conceptual framework depicted in Figure 1A. Here, the most hPTM and metabolite regulators with the highest edge weights are highlighted (**Table 1**), showing all modules they connect to, with red lines depicting anti-correlation and green lines positive correlations. The black pop-up shows the module properties and GSEA analyses of module 16. **Inset** ranks the modules according to the number of known CORUM complexes they encompass. Per module, enrichment for CORUM complexes was assessed using a hypergeometric test to determine whether the observed overlap exceeded what would be expected by chance. For each module, the bar chart displays the number of CORUM complexes that are significantly enriched. Bar height represents the count of significantly enriched complexes, while bar color indicates the combined p-value of these enrichments, calculated using Fisher’s method (e.g., if seven complexes are significantly enriched in module 3, their individual p-values are combined into one overall p-value). **B.** Module 16 is depicted by its individual protein expression patterns and its appointed hPTM and metabolite regulators, most prominently H3K27me3 and n6,n6,n6-trimethyllysine (Kme3), respectively. In orange, three different known protein complexes from CORUM are depicted, illustrating how the edges in the network can capture physically interacting proteins like the nuclear pore complex here. **C.** GSEA on module 16 as it is presented in the Lemonite report. D. Snippet from the Bigomics Correlation analysis tab, showing the highest correlating proteins over all 4566 proteins retained in the proteomics project. Here, pearson correlations can be easily inspected, e.g. to detect how one cell line like MOLM-16, highlighted in the legend, can drive certain PCC values as outliers. All these proteins associate in the nuclear pore complex, except for DNM2, which regulates microtubule dynamics (specifically acetylation) and the positioning of the microtubule-organizing center (MTOC) relative to the nucleus.

#### Protein Module composition

The modules in the network are clusters of proteins that consistently change over the 18 different AML cell lines, i.e. 18 different genotypes. This does not imply that these proteins physically interact or directly regulate each other’s expression. Yet, remarkably, the most correlating proteins effectively are part of the same protein complexes. In fact, 57% of the modules encompass (several different) known protein-protein interactions (PPI) or protein complexes enlisted in CORUM, including e.g. the Nuclear pore complex (NPC) in module 16 depicted in **Figure 3B and 3D** and Polycomb Repressive Complex 2 (PRC2), the NPC and the spliceosome A complex and E complex in Module 49. The inset in Figure 3A ranks the modules according to the number of CORUM complexes that are captured in each module ^25,26^. These are color-coded by how much these PPI enrichments deviate from what could be expected from random chance. Thus, the associations that bring together the proteins in these modules effectively can capture physical interactions in complexes, on top of mere functional interactions. In other words, molecular interactions in aptly complex experimental designs can be used as direct indicators of protein complexes without the application of interactomics workflows. In proteomics, these co-abundances across cell lines most probably reflect co-stabilisation within physical assemblies, i.e. protein complex dosage compensation.

### Targeting intra-omics edges: co-regulatory networks

The topology of the biomolecular network illustrates how these 18 different AML genotypes together allow defining consistently coherent biomolecular changes between omics layers, i.e. the edges can be quantified. Next, all the edges within each of the separate omics layers can be investigated from the original data matrix, comprising all quantified biomolecules. We first describe known connections that serve as confirmational foundation for moving towards more novel connections forward in future mechanistic follow-up studies.

#### Proteomics associations

For the proteomics data layer, all analytical perspectives are accessible as a public dataset in Omics Playground (Bigomics) (register in Omics Playground and open public dataset *AML_18CellLines_June2025*). This provides molecular biologists as well as medical doctors with browser-based interaction with the data through the full functionalities of the underlying R architecture with no need for coding skills.

CD34 is one of the most prominent biomarkers that defines AML cells’ stemness ^27^ and indeed CD34 is amongst the most discerning proteins in the data (**Supplementary Figure 4A**). While often associated to stemness, CD34 is a bit of a “sticky wicket”^28^. When approaching this data in a classic differential proteomics workflow, binary comparing CD34+ and CD34-cell phenotypes, a set of differential proteins emerges (n = 998, FDR < 0.01) (Figure 4A) with several known AML (ITGA6, ACSS1, MACROH2A2, RAB27A) and potentially novel biomarkers, including asparagine synthetase (ASNS). Considering the therapeutic impact of asparaginase treatment in ASNS^low^ acute lymphoblastic leukemia (ALL), depleting exogeneous asparaginase might be of benefit to evaluate asparaginase treatment in CD34+ AML^29^. Of note, in the Lemonite network, ASNS resides in module 31, which has Kme3 as its highest scoring metabolite regulator.

**Figure 4.**
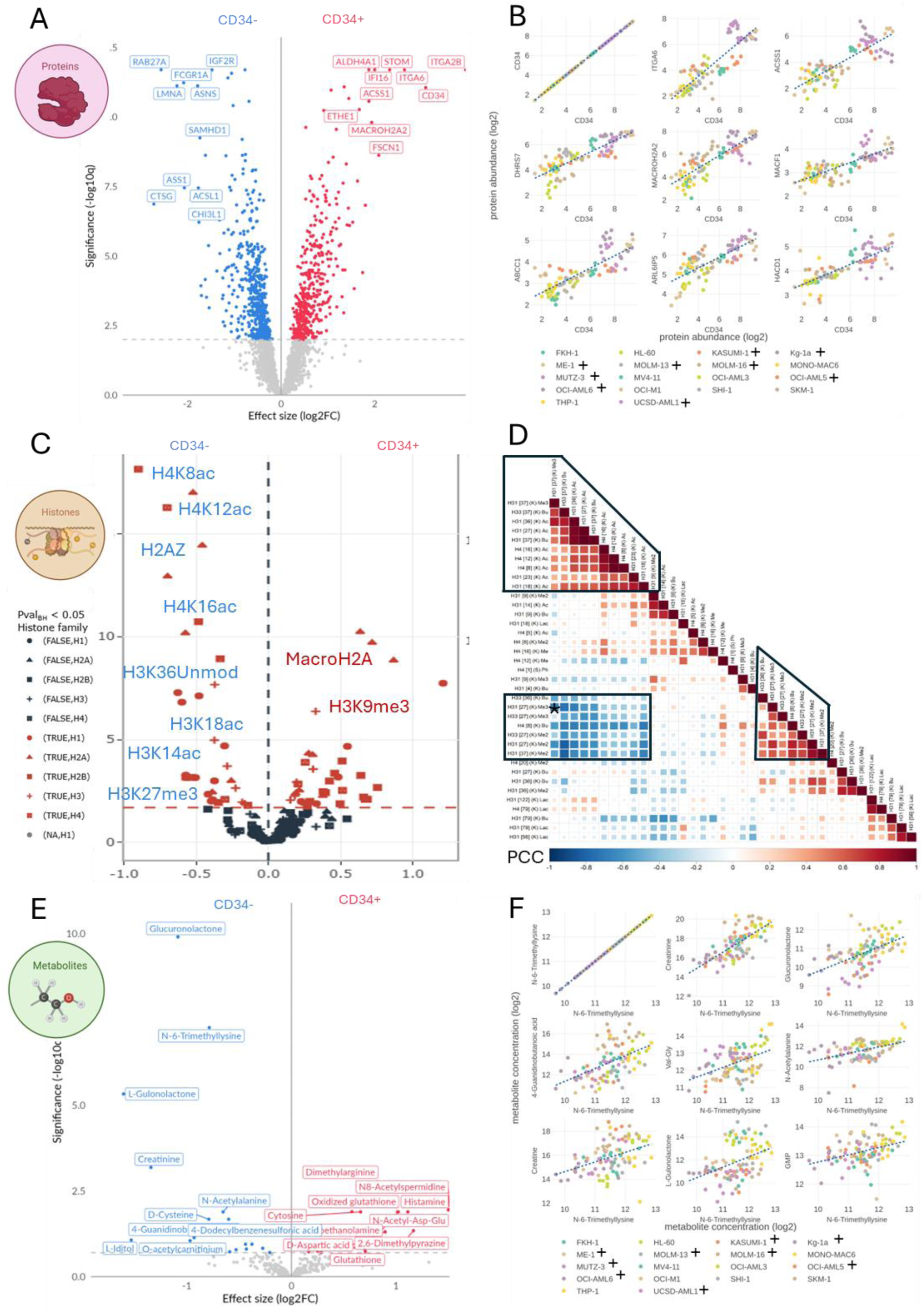
Targeting intra-omics edges. **A.** Volcano plot of differential proteins between CD34+ and CD34-AML cell lines. Several of the highlighted proteins were added to Figure 2. **B.** The 8 most correlating proteins to CD34 in the dataset (out of 4500+ proteins). Clearly, whenever CD34 expression increases, this also holds for these proteins. While this does not imply a direct causal connection, especially ITGA6 could very well be part of the same membrane-embeded complex with CD34. **C.** Volcano plot of the differential hPTMs between CD34+ and CD34-cells. Uniquely, we apply the msqrob2PTM statistical framework that was designed to robustly distinguish changes in PTM stoichiometry from changes in the abundance of the parent protein,i.e. Differential usage. PTM usage is then calculated through summarization, of all peptidoforms containing a given PTM. Finally, downstream differential analysis is done at both the peptidoform level and the PTM level using the well-established MSqRob2 framework. **D.** hPTMs can also be entangled. Certain hPTMs are signal integrators or hubs among hPTMs and consistently co- or anti-correlate. A correlogram depicting the binary Pearson correlation coefficients (PCCs) between all H3 and H4 hPTMs quantified. This highlights a clear positive correlation between most H3 and H4 acetylations (tip of the correlogram triangle) as well as an anti-correlation of these acetylations with H3K27me3 and me2, H3K36me and me2 and H4K8me/bu (monomethylations are depicted as butyryl in our workflow and are indistinguishable). **E.** Volcano plot of the differential metabolites between CD34+ and CD34-cells. The second most significantly differential metabolite is trimethyllysine, which derives from proteolysis of trimethylated proteins, including the very abundant histones. **F.** From the perspective of Kme3, two prominent pairs of the most positively correlating metabolites with an HMDB identifier in the Omics Playground (Bigomics) metabolomics project are creatine-creatinine and gulonolactone-glucurolactone. The first two are known to spontaneously convert, yet the latter co-occurrence raises a lot of questions, since they are not directly converted, while their co-regulation seems to argue that they do (Supplementary Figure 4K). Both are lactones of sugar acids and share similar cyclic structures, but glucuronolactone is involved in detoxification and connective tissue metabolism and gulonolactone is involved in vitamin C synthesis pathway, which cannot be completed in humans. Alternatively, one or both of the metabolite were wrongly annotated and manual curation provides another explanation.

Typically, such differential protein profiles are used for gene ontology analyses, most commonly using gene set enrichment analysis (GSEA). For the CD34 phenotype, the most prominent pathways found through GSEA are ***branched-chain amino acid catabolism*** (**Supplementary Figure 4B**) and ***Synergistic Anti-Leukemic Trc105 Decitabine GSE147503***. Branched-chain amino acid catabolism is increasingly recognized to underlie stem cell renewal capabilities in Eukaryotes, including haematopoietic stem cells ^30,31^ where HSC functional decline arises from redirecting BCAA usage from catabolic toward anabolic activity^30^. The decitabine GO term (a **DNA methyltransferase or DNMT inhibitor**) is a first line treatment in MDS, AML, JMML and CMML. This also implies that the use of cell lines does still allow to extract therapeutically relevant information. Of note, amongst the genes in the latter GO term are ITGA6, ITGA2B and STOM (**Supplementary Figure 4C**).

Importantly, there is no reason to assume that CD34 itself is inducing these functional shifts in AML cells. In fact, the edges help shortlist tightly correlating proteins that underlie more functional entanglements. Figure 4B visualizes the **8 most correlating proteins for CD34,** showing that any of these could be targeted just as efficiently to define CD34+/-phenotypes (Figure 2). **ITGA6** (Integrin alpha 6), has been identified as a marker for leukemia stem cells (LSCs) within the CD34+ population: long-term self-renewing LSCs reside within the CD34(+)/ITGA6(+) fraction ^32^. Next in line is ACSS1 (Acetyl-coenzyme A synthetase 2-like, mitochondrial), which catalyzes the synthesis of acetyl-CoA from short-chain fatty acids. It is therefore remarkable that on the other side of the volcano in Figure 4A is ACSL1 (Long-chain-fatty-acid--CoA ligase 1), which catalyzes the conversion of long-chain fatty acids to their active form acyl-CoAs. Thus, this implies that CD34 negative AML cells rely more heavily on the long chain fatty acids in their metabolism, in line with earlier findings ^33^ (Figure 2). This entanglement fundamentally impacts our understanding of biomarkers and is reflected prominently in more advanced biomarker discovery algorithms relying on machine learning. These attribute feature weights to proteins based on how much they help in clustering the different cell lines into the CD34-/+ phenotypes: they rank 15 proteins higher than CD34 itself in some iterations (**Supplementary Figure 4C**).

#### hPTM associations

Histones were processed in a new and more robust way visually explained in **Supplementary Figure 3^16^**. The comprehensive histone report of the variant-agnostic workflow can be downloaded from the Lemonite report page. The report includes a differential analysis on the co-extracted proteins (the acid extractome ^34^) as well as all three different data levels that can be inferred from the histone measurements, i.e. histone variants, peptidoforms (comprising combinatorial hPTMs) and single hPTM data. The “usage plots” that underly the single hPTM inference can be downloaded from this report to allow manual inspection of quantification (**Supplementary Figure 3C and 3D**). Of note, by lack of FDR control mechanisms in untargeted histone searching ^20^, manual curation of specific PTMs remains advisable. Therefore, the freely accessible Progenesis QIP project is available under **Supplementary Data 1** wherein other hPTMs can be searched, curated and quantified for subsequent analysis with the MS2Rob2PTM workflow. An extensive tutorial was published earlier ^35^. Here, we focus on single hPTMs only, which are presented as “differential usage”, as described elsewhere ^15^. Figure 4C displays the most differential hPTMs between CD34+ and CD34-cells as an exemplary contrast, showing the enrichment of acetylations in CD34-cells on both the H4 and H3 N-tail, with **H4K8ac** as the most upregulated hPTM. CD34+ cells are enriched for macro H2A (H2AY), as also seen in the proteome fraction (Figure 4A), and H3K9me3, amongst others. We also plotted the differential usage of hPTMs between the C-Mito and no-C-AML phenotypes defined elsewhere ^8^ (according to an in-house redefined clustering, **Supplementary Figure 4D**). Here, **H3K27me2/me3** is downregulated in the C-Mito cell lines, implying these cells have less of this repressive mark (**Supplementary Figure 4 E**).

To directly measure and visualize hPTM co-regulation (edges in the hPTM data layer), Pearson Correlation Coefficients (PCCs) between all hPTMs can be calculated and visualized in a correlogram (Figure 4D). For conciseness, only the H3 and H4 hPTMs are plotted (comprehensive hPTM correlations can be found in the html report of **Supplementary Data 3A**). This highlights a clear mutual positive correlations between most H3 and H4 acetylations as well as an **anti-correlation of these acetylations with H3K27me3 and me2, H3K36/37me and me2 and H4K8me**. We reported earlier that this anti-correlation between H3K27me3 and H4 N-tail acetylation is a hallmark in ESCs that are converting from naïve to primed state in both mouse and human ^4,5^.

#### Metabolite associations

In metabolomics, co-regulation of many metabolites is much more understood in terms of metabolic pathways. Thus, they are often a manifestation of changes in the proteome, whereby a set of enzymes is e.g. downregulated to shut down one pathway and activate another, because of changes in the energy availability in the environment (Figure 1A). As for hPTMs, no FDR thresholding exists and all targets of interest are best manually validated.

For the same CD34+/-contrast, the second most differential metabolite in the Omics Playground (Bigomics) project is **Kme3**, which is downregulated in CD34+ cells (Figure 4E). Amongst the most upregulated metabolites is dimethylarginine, another potential epigenetic echo in the metabolism, as it too is created by proteolysis of proteins carrying dimethylated arginines. When investigating the most correlated metabolites to Kme3 in the Omics Playground (Bigomics) project, two prominent pairs of positively correlating metabolites are emerging: creatine-creatinine and gulonolactone-glucurolactone (Figure 4F), which are also significantly downregulated in CD34+ cells (Figure 4E). The first two are known to spontaneously convert, yet the latter co-occurrence is more surprising, since they are not directly converted, other than the fact that they are lactone forms of sugar acids that sit in the same uronic acid pathway. In other words, the reason they correlate is most likely that both are passive overflow products of UDP-glucose and UDP-glucuronate metabolism. Examples like these serve as lead generators for future mechanistic follow-up studies.

### Functional Entanglement in the full cellular context

Building on the rich functional annotations present in the proteomics data layer, we further extended on **functional entanglement in the full cellular context.** More specifically, the Lemonite network relied exclusively on the most variable biomolecules and connected specific regulators in a competitive algorithmic approach to extract functional interpretations by GSEA on individual modules. However, for every biomolecule in the data, PCCs can be calculated over the 97 samples to all 4612 proteins, quantifying every edge to a protein in the data. For a given target, all the 4612 PCCs can then be ranked to capture what proteins are consistently co- or anti-correlating to this target. In turn, this can be made into an input for a GSEA analysis, providing functional entanglement in the full cellular context. These are browsable at the bottom of the Lemonite report under “Functional entanglement in the full cellular context**”** and PCC rank plots are available from **Supplementary Data 4**. This is akin to the conventional fold change GSEA analysis in a binary comparison between two cellular phenotypes, but this time the PCCs are used to study the functional environment of a single biomolecule. We illustrate this concept consecutively on protein functional entanglement, hPTM functional entanglement and metabolite functional entanglement. So, while individual protein functions are often derived from targeted intervention studies and loss-of-function approaches, our experimental design provides the functional context for all the biomolecules detected in undisturbed phenotypes, which is not the single biomolecular function itself.

#### Protein functional entanglement: CD34

As demonstrated in Figure 4B, many proteins have strongly correlated expression profiles with functional implications for e.g. CD34. On the other side of the spectrum, many are strongly anti-correlating (including LMNA, EPRS1 and ASNS). A GSEA on the ranked PCCs to all proteins therefore provides a comprehensive picture of what functionally changes in a cell where CD34 is up- or downregulated, hereby not implying that CD34 is causing the changes directly. Most prominently, CD34 is positively associated with *branched-chain amino acid metabolism* (as for the binary fold-change comparison) and *hematopoietic stem cell differentiation*, nicely demonstrating the power of this functional entanglement approach, given that CD34 is a stemness marker in AML (Figure 5A and B). In fact, Branched-chain amino acid metabolism has been known to improve age-related reproduction in *C. elegans* and more recently was found to restore replication-dependent hematopoietic stem cell fitness in haematopietic stem cells ^30,31^. Additionally, CD34 is negatively associated with *cytoplasmic ribosomal proteins, Aminoacyl tRNA biosynthesis, proteasome degradation* and *Orc1 removal from chromatin* and positively to *shingolipid metabolism* and *O-acyltransferase activity* (fundamental to lipid metabolism) (**Supplementary Figure 5B**). In a sense, when CD34 expression is approached as a biomarker, these functions in AML cell lines can be expected to shift with it (Figure 2).

**Figure 5.**
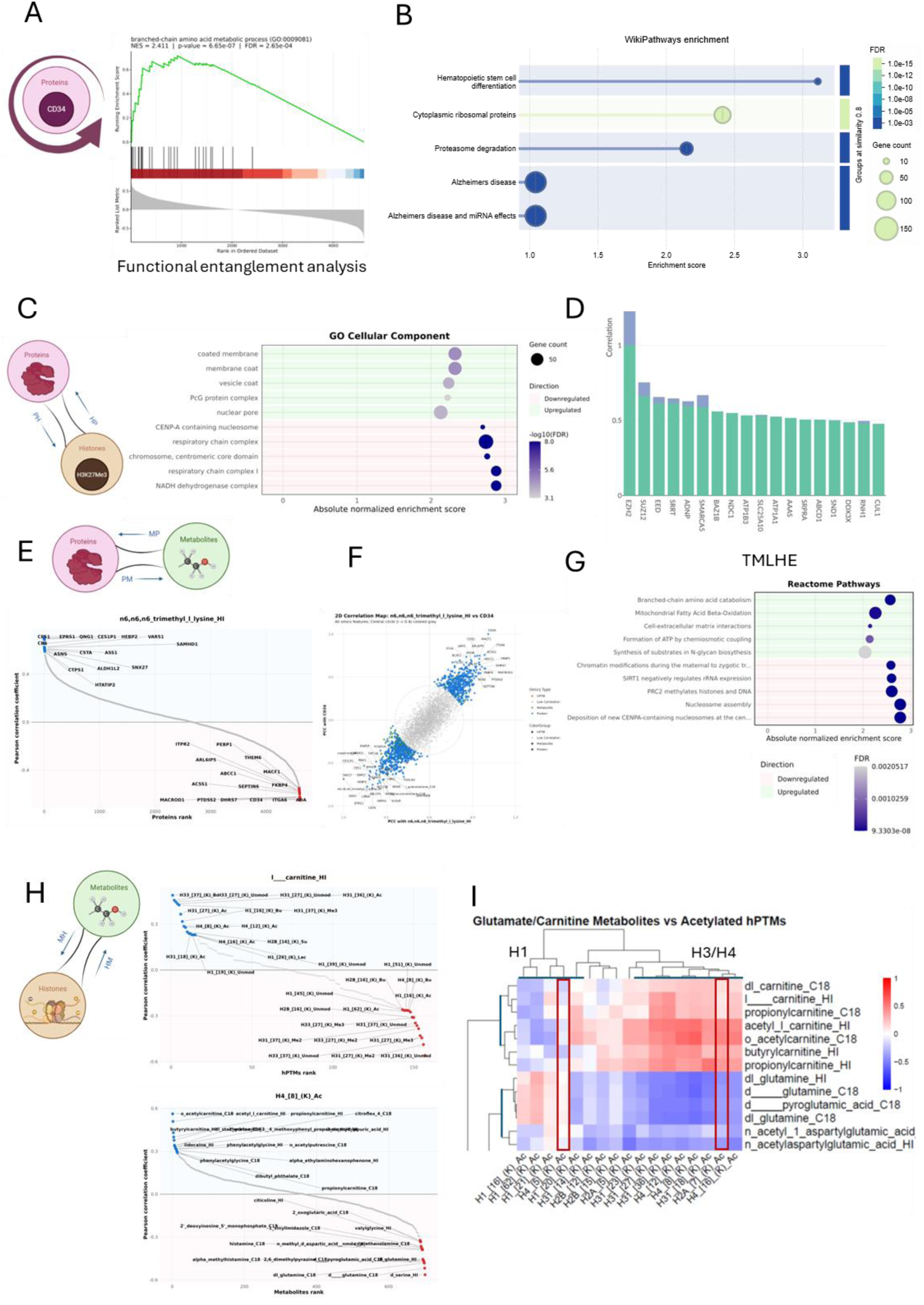
Functional entanglement analysis. As opposed to comparing cellular phenotypes by performing GSEA analyses on ranked protein fold changes from a given contrast, ranking all PCC values to all proteins in the design, provides a functional entanglement analysis that surfaces coherent biomolecular behaviour within the experimental design. Two different PCC rankings are available: one derived from Omics Playground (Bigomics) (which is built directly on the DIANN output) and one from the MSqRob protein inference report (used to build the Lemonite network). **A. Protein-protein functional entanglement.** The ranking was used to study CD34 first, showing that - as for the binary cellular contrast - CD34 associates with *branched chain amino acid metabolism.* **B.** When searched in STRING db, the highest scoring pathway under Wikipathway (enrichment score 3.10 and FDR 0.00068 in STRING) was Haematopietic Stem cell differentiation, albeit with “both ends” under direction, with RIOK3 and TRAF3IP3 displaying anti-correlation. **C. hPTM-protein functional entanglement.** A functional entanglement analysis on H3K27me3, building on all protein PCCs for H3K7me3 (HP/PH axis in Figure 1A). Several of the GO terms were also detected in different modules in the Lemonite biomolecular network for H3K27me3, including *nuclear pore* and *PcG complex* and a negative enrichment of *mitochondrial respiratory chain complex*. One interesting GO term is the anti-correlation to CENP-A/centrosome. CENP-A is the histone H3 variant that replaces canonical H3 nucleosomes at the centromere to act as the essential, epigenetic foundation for kinetochore assembly (not searched in our hPTM analysis). They function as a "pedestal" to recruit specific proteins like CENP-C and CENP-N. When targeting CENP-C we found that its 10th highest ranking protein partner is RBBP4 (of the PRC2 complex) and the third highest is in fact KDM2A in Omics Playground (Bigomics), which demethylases H3K36me3. In **figure 4D**, H3K37me3 (indistinguishable from K36 by mass spectrometry) strongly anti-correlates with H3K27me3, an anti-correlation studied extensively elsewhere (ref). **D.** The most correlating proteins to EZH2, out of 4612 are two other members of the PRC2 complex (EED and SUZ12) as well as SMARCA5 and BAZ1B. Therefore, SRRT, part of the primary microRNAs (miRNAs) processing machinery during cell proliferation, as well as ADNP most probably also associates with the RPC2 complex. **E. Metabololite-protein functional entanglement.** Ranked correlations of all proteins to Kme3. The most anti-correlating proteins are CD34 and its entangled proteins. **F.** 2D-plot of the protein correlations to all biomolecules for Kme3 and CD34, illustrating how they are at two ends of the biomolecular network. **G.** Functional entanglement of TMHLE that converts Kme3 into carnitine shows its strong anti-correlation it to the PRC2 complex, the main mediator of H3K27me3, as well as a strong correlation to branched-chain amino acids, which also the most prominent functional entanglement of CD34, shown in A. **H. The Energy-information axis, i.e. hPTM-metabolite entanglement.** The highest ranking hPTMs to carnitine are acetylated H3 and H4 residues. Inversely, H4K8ac correlates most strongly to all acyl-carnitines (depicted in **Supplementary Figure 5C**). Most strongly anti-correlating are glutamine and serine. **I.** A heatmap depicting several glutamine and carnitine derivatives and the acetylations in the data. These two classes of metabolites cluster the different histone acetylation nicely, with H3 and H4 N-tail forming a dense cluster.

#### hPTM Functional entanglement: H3K27me3 and H4K8ac

As for proteomics, the calculated PCCs of an hPTM to all 4612 proteins can be ranked to become an input for a GSEA analysis that provides inter-omics functional entanglements. While the values overall are lower than protein-protein PCCs, http://www.lemonite.ugent.be/AML_Results provides a total of 158 hPTM functional entanglements combining into a Functional Entanglement Atlas for the histone epigenome of AML. **Supplementary Data 4** provides all these calculated PCC values to also allow easy input into e.g. STRING-db for extended analysis against e.g. InterPRo, SMART and other GO classes (www.string-db.org).

Here, we again focus on the most tightly connected hPTMs in the biomolecular network, i.e. H3K27me3 and H4K8ac. Performing a GSEA on H3K27me3/protein PCCs confirms the earlier finding from the biomolecular network from every GO perspective, with a positive enrichment of *PcG Protein complexes* and a negative enrichment of *mitochondrial respiration* and *CENPA-containing nucleosomes* (Figure 5C). An additional GO term is the positive correlation between H3K27me3 and *nuclear pore (NCP) organization*, which had also been captured in the Lemonite network in module 16 (Figure 3B). Unknown until recently, Nup93 has now been found to co-localize with Polycomb (PRC2) proteins and H3K27me3-marked chromatin regions in Drosophila ^36^. Indeed, in the multi-omics Omics Playground (Bigomics) project, EED ranks as the first protein out of all 5521 multi-omics feature correlations to H3K27me3, proving the applicability of inter-omics functional entanglement studies. In other words, the **PH directionality in** Figure 1A can be directly deduced from the data, i.e. the writer complex defines the H3K27me3 abundance. Unintuitively however, **MAT2A** is the 7^th^ highest ranking anti-correlating protein to H3K27me3, which imply lower SAM availability in H3K27me3^High^ cell lines. **Propionyl-carnitine** is the most anti-correlating metabolite in this multi-omics correlation list, confirming the above findings of module 21 from the lemonite network.

Protein complexes can be reconstructed from the PCC values (Figure 3). Ranking all protein correlations to the other PRC2 member **EZH2** shows just how noiseless the data is: the two highest correlating proteins out of 4566 entries retained in the proteomics Omics Playground (Bigomics) project, are **SUZ12** and **EED** (Figure 5D). This, in fact, turns out to be a more universal principle in proteomics data of more complex experimental designs: when loading the data of Kramer et al. on patient bone marrow AML samples into Omics Playground (Bigomics) ^37^, the same phenomenon could be reconstructed, whereby the 1^st^ and 2^nd^ most correlating proteins to EED out of 7771 are SUZ12 and EZH2 and 6 proteins in the top ten are nucleoporins (**Supplementary Figure 5B**). In turn, this implies that the other highest-ranking proteins to EZH2 are also functionally entangled to H3K27me3, or even part of the same complex. **SMCA5** (SWI/SNF-related matrix-associated actin-dependent regulator of chromatin subfamily A member 5), an ATPase that possesses nucleosome-remodeling activity, indeed was identified as the 5^th^ highest ranking protein. It is an essential component of the NoRC (Nucleolar Remodeling Complex), together with the regulatory subunit **BAZ2A** (bromodomain adjacent to zinc finger domain 2A), ranked 111^th^ (Figure 2). **BAZ1A** ranks 44^th^, yet more strikingly, another bromodomain adjacent to zinc finger domain, **BAZ1B**, effectively ranks 6^th^ in the PCC list of EZH2, just after SMCA5. Excitingly, a recent study on the PRC2 interactome in which we were involved, found BAZ1B as one of the PRC2 interactors in a BioID study ^38^. Inversely, when a functional entanglement was performed on BAZ1B, *PRC2 methylates histones and DNA* lists as its first hit in Reactome (FDR: 3.3e-8). Other members correlating positively include **RBBP 4 and 7** and **MTF2** (Metal-response element-binding transcription factor 2). MTF2 is also the 3^rd^ highest ranking protein correlating to **EED** in the data of Kramer et al.^37^ (**Supplementary Figure 5B**). MTF2 reinforces the PRC2.2 specifically, which is the variant most associated with CpG island targeting and maintenance of repression at developmental loci — exactly the genes that need to stay silent to preserve the AML differentiation block.

At first glance less obviously, the 3^rd^ and 4^th^ proteins in Figure 5D, are **SRRT** and **ADNP**. **ADNP** (activity-dependent neuroprotective protein) is the more interesting case based on prior knowledge, so SRRT is the more interesting for mechanistic follow-up. ADNP is the defining subunit of the ChAHP complex, which contains ADNP together with HP1 proteins and the chromatin remodeller CHD4 (which also correlates positively). ChAHP is a repressive complex that operates at a partially overlapping but distinct set of genomic targets from PRC2. Most probably, RBBP4/7 form the key bridge as these two histone-binding WD40 proteins are genuinely promiscuous — they appear as shared subunits in PRC2, NuRD, CAF-1, and the NURF/ISWI complexes. In proteomic abundance data, a protein that is shared across multiple complexes will correlate with all of them simultaneously, even if those complexes never directly interact. It is clear then, that changing the perspective now to the functional entanglements of RBBP4 will deepen our understanding of the repressive machinery activated in AML.

One final, striking positive correlation is that with **HDAC2** (Histone deacetylase 2). Excitingly, Figure 4D already captured an anti-correlation in the histone layer that implied the involvement of an HDAC protein: H4 N-tail acetylation is lower whenever H3K27me3 is higher. This anti-correlation was already described by us in human and mouse ESCs, but hitherto has no direct mechanistic description ^4,5^. In turn, this shows what the edges in this dataset represent: they are biomolecular “behavioral” correlations that also capture direct interactions. Each of these extra order associations can become the point of perspective for a next analysis, thus gradually changing the perspective on the network. Here, we started from H3K27me3 to find its writer complex, PRC2, which is associated with the nuclear pore complex, the NoRC complex and other interactors like BAZ1B and HDAC2, the latter of which capturing the anti-correlation with abundance of H4 N-tail acetylation found in the hPTM omics layer (Figure 4D).

#### Metabolite functional entanglement: trimethyllysine and carnitine

From the Lemonite biomolecular network **Kme3** was found to be a densely connected metabolite which is released from cognate proteins like histones via proteolysis. It serves as a precursor for carnitine biosynthesis. Strikingly, when ranking all 4566 protein PCCs to Kme3, we directly sample the **MP/PM edge in** Figure 1A and the most anti-correlating protein is **CD34** (PCC: -0.685), with evidently **ITGA6, ACSS1** and **ABCC1** equally part of the top 10 highest correlating proteins (Figure 5E)**. MACROD1**, just below CD34, deacetylates O-acetyl-ADP ribose, a signalling molecule generated by the deacetylation of acetylated lysine residues in histones and other proteins ^39^. In the top 20 most positively correlating proteins to Kme3 are again **ASNS**, **VARS1**, **ASS1**, **LMNA** and **EPRS1**, other biomarkers in CD34+ AML cell biology (Figure 2). On a 2-dimensional dot plot depicting PCC values for Kme3 and CD34 to all other biomolecules in the matrix, this deep anti-correlation is captured even more elaborately (Figure 5F).

From another perspective, Mitochondrial 6-N-trimethyllysine dioxygenase epsilon (TMLHE) is the mitochondrial enzyme converting Kme3 into 3-hydroxy-6-N-trimethyllysine as the first step in carnitine biosynthesis. A functional entanglement analysis of the TMLH protein strongly anti-correlates to the PRC2 complex and positively to branched chain amino acid metabolism (Figure 5G). Counterintuitively therefore, there is lower expression of TMLH in cell lines with higher PRC2 expression.

### The energy-information axis

Finally, we explore the energy-information axis directly. With little direct correlation between Kme3 and PRC2, i.e. H3K27me3, we investigate the carnitines instead. Indeed, carnitine, acetyl-carnitine and propionylcarnitine, all correlate positively to many of the acetylation events shown in Figure 4D, therefore anticorrelating to e.g. H3K27me2/3 (Figure 5H and **Supplementary Figure 5C**).

Inversely, when considering the correlations of all metabolites to H4K8ac, all the acyl—carnitines rank high. Here it is worth revisiting the Lemonite network, where the mTOR-containing module 72 connects H4K8ac and carnitine directly. This small module also contains other metabolic hints, including COX7A, component of the cytochrome c oxidase (OxPhos) and NMNAT1, which catalyzes the formation of NAD^+^ from nicotinamide mononucleotide (NMN). Additionally, the module has a strong transcriptional component, comprising spliceosome factors PRPF38A, PRPF6, SMARCE1 and SNRNP40 along INTS3, HMGB1 and GTF2H4. Surprisingly, CTR9 also strongly correlates with mTOR and is a component of the PAF1C complex, which is involved in hematopoiesis and stimulates transcriptional activity of KMT2A/MLL1.

On the other side of this spectrum, two detections of glutamine that were depicted in from Figure 3A are the most anti-correlating metabolites, together with serine (Figure 5H). A heatmap of the different glutamine and carnitine metabolites plotted against all (?) detected acetylations, finally show that this inverse correlation is most prominent for histone H3 and H4 (Figure 5I).

## DISCUSSION

The genome provides an organism with a set of instructions to make building blocks that can be rearranged into a very rich combinatorial space that dictates different cellular phenotypes. This could be referred to as the cellular “phenospace” of a given genome. This is a product of the cellular history, which in turn defines and is defined by the environmental context in which the cell is localized at a given time of interest. Importantly, the phenospace in not a continuum of all potential molecular combinations. Rather the options for a cellular phenotype are a grained manifestation of the manifold interpretations of a given genome. In the language of dynamical systems, these are called attractor states or attractors for short. Attractors are the modern equivalent of Waddington’s valleys ^1^, which seem to be a general property of complex, multicellular organisms: small changes push the system only slightly away from a stable state (imagine rolling Waddington’s ball a little up the valley wall) and it will then return to its attractor state. Only when larger/functional/co-expressional changes occur, does the cell “roll” into another attractor state, in turn inducing other co-ordinated biomolecular changes. This is intuitively true in a developmental context, because we know that there is only a specific subset of cell types and tissues that together make up a healthy human body. One such example are the different myeloid cell types. Yet in cancer, “new” cellular phenotypes are being explored that no longer obey the checks and balances of normal development and therefore overgrow the healthy organism, such as is the case in AML. Inversely, the phenotypic manifestation of a given myeloid cell type, or a cancerous derivative thereof, also is a product of its genotype and cellular history yet is equally coarse-grained by the fact that molecules cannot change their expression in isolation, i.e. without altering that of other biomolecules. It is by presenting 18 different AML genotypes that we capture the latter granularity: the biomolecules that behave coherently, do so independent of the genotype and their behaviour in not encoded in the genome. In turn, these patterns are crucial to predict or study drug efficacy and adverse outcomes, independent of patient genomes, particularly in a cancer caused by differentiation arrest.

Because the biomolecular network contains over 5500 proteins, hPTMs and metabolites, close the 30 million edges can be calculated. We therefore make this data accessible to the broader community, with this manuscript illustrating its use. We propose the term functional entanglement to contrast our direct multi-omics measurement in a single experiment with more overarching and public data efforts in systems biology ^10^. We define three classes of functional entanglement: (i) most tightly correlated proteins often reside together in complexes, as shown by CORUM enrichment of the protein modules as well as the top-ranking proteins in the full PCC protein rankings in our and in other data^37^. We propose to extend the term protein complex dosage compensation ^17^ and show how this direct measurement provides hitherto unknown binding partners for follow-up studies. Indeed, this data even allows defining inter-omics substrate binding, as exemplified by the strong correlation in abundance between H4K8ac and its reader protein BRD4, which seems to imply mutual stabilization. (ii) Proteins that are functionally correlated through the biomolecular perspective presented in Figure 1A. Here, known interactors serve as positive controls for the overall value of the network and the interspersed proteins in protein modules serve as leads for mechanistic follow-up studies. (iii) Functionally entangled biomolecules that have no obvious causal relationship. The latter effectively surfaces the very relativity of causality: how many consecutive causal interactions are still considered a causal interaction between the two ends of the chain? Our short exercise herein loops back onto itself: H3K27me3 correlates strongly to its PRC2 proteins, which themselves correlate to other (recently discovered) interactors, including HDAC2, which in turn could underlie the known anti-correlation between H3K27me3 and H4 N-tail acetylation which we also described earlier ^5,38^. Yet, cascading through n^th^ order correlations in this data matrix gradually turns around the perspective on rather than simply providing increasing numbers of equally correlating biomolecules. Indeed, despite the lack of edge directionality between single nodes, it is important to realize that there is directionality in a one-vs-many edge: the ranking of correlations differs depending on the starting molecule chosen. Indeed, when a partner protein ranks first from the perspective of a given protein, the reverse perspective often shows a set of proteins correlating even more to the partner protein, like its protein complex partners. As such, at one point these functional correlations are potentially no longer causal, or this at least becomes a more philosophical question. Yet, since they are persistently observed to be present, we denote them as a new and third class of functional entanglements.

All these classes of correlation have known equivalents in systems biology, but this is intrinsically different as we have measured them directly in a single experiment, and between hitherto unexplored biomolecular layers extracted from single cell pellets. To frame the relationship between the different omics, we also propose a circular biomolecular perspective wherein e.g. epigenetic changes in histones lead to newly expressed proteins that convert given metabolites, which are projected back onto the histones in the form of hPTMs. This completes a feedback loop—an ‘energy-information’ axis—where histones serve as the signal integrators, regulating gene activity based on the metabolic state of the cell. This view goes back to the first Eukaryote merger, where the Archaeal contributor provided the histones, allowing genome expansion and better control of gene activity by reading out the energy state of the bacterial endosymbiont that became the mitochondrion^11^. To date, the Eukaryote cell still senses this cellular energy state and metabolic flux through histones, adjusting protein expression accordingly ^40^. In cancer biology, altered metabolism is not merely a byproduct of tumorigenesis but rather a requirement for tumor initiation and progression ^41^. In this new view, hPTMs become - at least in part - dysregulated due to metabolic changes, contributing to uncontrolled cell proliferation, impaired differentiation, and other hallmarks of malignancy through altered protein expression. From the data perspective, then, the class of functional entanglement becomes a function of the number of loops required in the circular graph to explain the observed correlations.

The applicability of this circular diagram is illustrated by the fact that it implied the existence of an unobvious edge directionality, i.e. the histone-metabolome connection (**HM** in Figure 1A). We found at least two such examples in the data: the metabolite trimethyllysine (Kme3) and dimethylarginine, which can only derive from proteolysis of proteins that carry these methylations as a PTM. In the Lemonite network, this metabolite connects to the same module 16 as does H3K27me3, yet metabolic tracing would be required to show a direct link with this specific hPTM. Thus, this biomolecule, Kme3, potentially goes through a drastic functional transition, going from an epigenetic signal integrator within the histone H3 backbone, to a metabolite that is actively converted into acyl carnitines that mediate lipid metabolism and even might drive H4 N-tail acetylation further downstream. It is such connections that underlies e.g. the inverse correlation of the converting enzyme THMLE with PRC2 complex in functional entanglement analysis on the ranked PCCs. This is an exciting example not only of less intuitive directionality, but also of the emergent nature of biomolecular function in general ^1^.

Importantly, measurement of these three different omics layers yields (inferred) abundances of the analytes but not their biological activity, which complicates the direct functional interpretation of the edges. First, if a protein is no longer functional because it has a mutation, its abundance is of little relevance, as illustrated in our **Supplementary Figure 1**. Therefore, genomics remains the fundamental layer on which these phenotypic multi-omics analyses are built. Second, lack of the required metabolite co-factors can render the protein inactive. Here, the read-out of the measured metabolites could provide indirect telltale insights into enzyme activity. Third, the absence of another protein in the complex could render the complex inactive, as exemplified in **Supplementary Figure 1** and capturing the abundance of the full complex – captured in the most tightly correlated proteins – can help resolve this. Also, PCC as metric is very sensitive to outliers and requires manual verification before extending the analysis. For example, THP-1 and UCSMDI-AML cluster away from all other cell lines in the metabolome UMAP (Figure 1C).

In conclusion, the power of this novel multi-omics approach derives from (i) the unique combination of the three functionally entangled phenotypic omics layers, i.e. the hPTMs, the metabolome and the proteome; (ii) the fact that all three omics layers are isolated sequentially from the same cell pellets creates deep resolution between the cellular phenotypes and unprecedented coherence between the omics layers; (iii) the hPTM abundances are inferred within a new robust statistical framework; (iv) the adequately complex experimental design of close to 100 samples, comprising 18 different AML cell lines, i.e. phenotypes, allows robustly quantifying correlations within and between omics layers and allows detection of novel functional entanglements, including novel protein complex members, (v) measuring the unperturbed cell states, i.e. the phenotype that is established through different genetics and cellular history, captures the emergent yet consistent patterns in the biomolecular phenospace and (vi) most importantly, all the data can be manually browsed and curated without bioinformatics skills. We therefore also provide a video tutorial to help surface all the hundreds or thousands of edges still hidden in this dataset. We foresee a future where similarly rich multi-omics datasets are made available in the same way, so that these become a much-needed hypothesis repository for labs that do not have direct access to advanced mass spectrometry platforms that can acquire such elaborate sample batches, further democratizing MS-based omics. After studying this network intensely, it almost seems to evolve from descriptive to mechanistic. The well over 30 million values that can be attributed to the edges become a reflection of the biomolecular interactions that underlie Eukaryote complexity. So, if the complexity of the network and data structure feels overwhelming, that is because biology is overwhelmingly complex.

## METHODS

### 1. Cell culture

Acute myeloid leukemia cell lines (n=18) were obtained from the German Collection of Microorganisms and Cell Cultures GmbH (DMSZ) and cultured according to manufacturer’ s guidelines (supplementary Table). Culture media was supplemented with 10% or 20% heat-inactivated fetal bovine serum (Gibco™) (included in the metadata in the Omics Playground (Bigomics) Project), 1mM sodium pyruvate (Gibco™), 2mM Lglutamine (Gibco™), penicillin (100U/ml)-streptomycin (100 μg/ml) and amphotericin B (0.25 μg/ml)(Gibco™). Cells were maintained under 5% CO2 at 37°C. Cell cultures were verified to be free of mycoplasma using the MycoAlert® Mycoplasma Detection Kit (Westburg Life Sciences, Inc.). Cells (1x106) were harvested by centrifugation (5min, 1500 RPM, 4°C) and washed with cold PBS (1X) (Gibco™). The resulting cell pellets were immediately snap-frozen in liquid nitrogen and stored at -80°C until further processing. For each cell line, six biological replicates were harvested at distinct time points to account for biological variability.

### 2. Histone sample preparation

Histone extraction of the cells was performed using direct acid extraction as optimized and described previously ^21,42^. Additionally, in this workflow, the pelleted fraction after HCl addition (containing the proteome) and the supernatant after TCA precipitation (containing metabolites) were preserved for subsequent sample preparation. Consequently, one-dimensional SDS-PAGE on a 9-18% TGX gel (Bio-Rad Laboratories) was performed on a fraction of the resulting histone extracts for quantification and normalization ^43^. The remaining histone extracts were propionylated and digested following an optimized protocol ^18,19,44^.

### 3. Metabolome sample preparation

The metabolomics was performed by BGI genomics. Internal standards were added: d3-Leucine, 13C9-Phenylalanine, d5-Tryptophan, 13C3-Progesterone. Metabolite extraction was done as follows: 25 mg sample was weighed, added into 2 ml thickened centrifuge tubes, and 2 magnetic beads were added. 10 μL of the prepared internal standard 1 was added to each sample. 800 μL of precooled extraction reagent was added (methanol: acetonitrile: water (2:2:1, v/v/v)), and samples were ground at 50HZ for 5min. The ground samples were placed at -20°C for 2h and the sample were centrifuged (25000 g *4℃, 15min). 600μL of each sample was added in split-new EP tubes, then freeze dried. 120 μL of 50% methanol was added to the dried sample and shaken until completely dissolved. This was centrifuged at 25000 g *4℃ for 15min. The supernatant was aspirated and placed it in a split-new EP tube. 10 μL of each sample was mixed into QC samples.

### 3. Proteome sample preparation

Pelleted proteins, following histone extraction, were digested using the S-Trap protocol (ProtiFi). Lysis buffer (10% SDS, 100 mM triethylammonium bicarbonate, TEAB) was added to the liquid sample in a 1:1 ratio. The proteins were solubilized by sonication and vortexing. Reduction was performed by adding dithiothreitol (DTT) to a final concentration of 5 mM, followed by incubation at 37°C for 30 min at 600 rpm in a ThermoMixer Comfort (Eppendorf). Alkylation was carried out by adding iodoacetamide to a final concentration of 15 mM, with samples incubated in the dark at room temperature for 30 min. To acidify the proteins, 12% phosphoric acid was added. Binding/wash buffer (100 mM TEAB in 90% methanol) was then introduced, resulting in a fine protein suspension. The suspension was loaded onto S-Trap columns (ProtiFi) and centrifuged at 4000 × g for 30 s in a refrigerated centrifuge (Eppendorf 5417R) to trap the proteins in the depth filtration plugs. The trapped proteins were washed three times with binding/wash buffer, followed by centrifugation at 4000 × g for 30 s after each wash. Proteins were digested directly on the S-Trap columns by adding a digestion buffer containing 1 μg trypsin/lys-C per sample. Digestion was carried out at 47°C for 3 hours in a thermoshaker. Peptides were eluted sequentially with 50 mM TEAB, 0.1% formic acid (FA), and 50% acetonitrile in Milli-Q water, with centrifugation at 4000 × g for 1 min between each step. The eluted peptides were dried using an SPD111V SpeedVac concentrator (Thermo Scientific) at 50° C, and the dry samples were stored at -20°C until further analysis.

### 4. Liquid chromatography and mass spectrometry analysis

Histone and proteome samples were resuspended in 0.1% formic acid and injection volumes were adjusted resulting in 800 ng of histones and 200 ng of peptides on column, respectively. A quality control mixture was created by mixing 2 μl of each sample. Trapping and separation of the peptides was carried out on a Triart C18 column (5 mm × 0.5 mm; YMC) and a Phenomenex Luna Omega Polar C18 column at 45°C (150 mm × 0.3 mm, particle size 3 μm), respectively, using a low pH reverse-phase gradient. Buffers A and B of the mobile phase consisted of 0.1% formic acid in water and in acetonitrile, respectively. A 20-min non-linear gradient was used going from 2% to 50% Buffer B, followed by a washing step at 90% mobile phase B and an equilibration step at 2% mobile phase B (starting conditions). The samples were run in a randomized fashion and a quality control injection of a mixture of all samples in the experiment was incorporated every ten samples. Data acquisition was performed on a ZenoTOF 7600 system (AB Sciex) operating in positive mode coupled to an ACQUITY UPLC M-Class System (Waters) operating in capillary flow mode (5 μl min−1).

For histone analysis, samples were analyzed using data-dependent acquisition (DDA) mode. Each cycle consisted of a full MS1 scan (m/z 350–1,250) with a duration of 100 ms, followed by an MS2 scan (m/z 140–1,800, high-sensitivity mode) with a duration of 12 ms. A maximum of 40 precursors (charge state +2 to +6) exceeding 1500 c.p.s. were monitored, followed by an exclusion for 3 s per cycle. A collision energy of 12 V and a cycle time of 800 ms was applied. Ion source parameters were set to 4.5 kV for the ion spray voltage, 35 psi for the curtain gas, 15 psi for nebulizer gas (ion source gas 1), 60 psi for heater gas (ion source gas 2) and 200 °C as source temperature. Proteome samples were analyzed using data-independent acquisition (DIA) mode with a Zeno SWATH acquisition scheme. The setup included 85 windows spanning a precursor ion mass range of 399.5–903.5 Da, employing dynamic collision energy with CID as the fragmentation mode and an accumulation time of 13 ms. Ion source gas 1 and 2 were set to 20 and 55 psi respectively and curtain gas was set to 45 psi (80). MS1 covered a mass range of 400 – 1200 Da with an accumulation time of 0.1 s and had a collision energy of 12 V. MS2 covered a mass range of 140 – 1800 Da.

For UPLC-MS Analysis, Waters UPLC I-Class Plus, Waters, USA, was coupled to a Q Exactive high resolution mass spectrometer (Thermo Fisher Scientific, USA). Chromatographic separation was performed on a Waters ACQUITY UPLC BEH C18 column (1.7 μm, 2.1 mm × 100 mm, Waters, USA), and the column temperature was maintained at 45 °C. The mobile phase consisted of 0.1% formic acid (A) and acetonitrile (B) in the positive mode, and in the negative mode, the mobile phase consisted of 10 mM ammonium formate (A) and acetonitrile (B). The gradient conditions were as follows: 0-1 min, 2% B; 1-9 min, 2%-98% B; 9-12 min, 98% B; 12-12.1 min, 98% B to 2% B; and 12.1-15min, 2% B. The flow rate was 0.35 mL/min and the injection volume was 5 μL. Q Exactive (Thermo Fisher Scientific, USA) performed the primary and secondary mass spectrometry data acquisition. The full scan range was 70‒1050 m/z with a resolution of 70000, and the automatic gain control (AGC) target for MS acquisitions was set to 3e6 with a maximum ion injection time of 100 ms. Top 3 precursors were selected for subsequent MSMS fragmentation with a maximum ion injection time of 50 ms and resolution of 17500, the AGC was 1e5. The stepped normalized collision energy was set to 20, 40 and 60 eV. ESI parameters: Sheath gas flow rate was 40, Aux gas flow rate was 10, positive-ion mode Spray voltage(|KV|) was3.80, negative-ion mode Spray voltage(|KV|) was 3.20, Capillary temperature was 320°C, Aux gas heater temperature was 350°C.

### 5. Proteome data processing

Analysis of proteomics DIA data was performed using DIA-NN (version 18.1)^45^. Raw files were imported into DIA-NN, and preprocessing steps included noise reduction and peak picking. A spectral library was generated from a FASTA file containing human protein sequences retrieved from Uniprot (accessed on Dec 6th, 2023), supplemented with contaminants. Peptide identification was performed using a combination of library matching and deep neural network prediction using double-pass mode as neural network classifier. Quantification was performed using the robust LC high accuracy setting. Mass accuracy was set to 20 ppm and MS1 accuracy was set to 10 ppm.

Protein quantification was done through two parallel tracks: one leading to Omics Playground (Bigomics) and one to Lemonite. For the Bigomic input file, the protein abundance table from DIA-NN was exported, allowing single peptide annotations (requiring retrospective manual verification in e.g. Skyline for biomarkers). Only samples were retained that were also retained in the histone and acid e xtractome fractions (Lemontree). Imputation was done with SVD, Keep NA rows was checked with >= 3 in any group, median normalization, no outlier removal, no batch correction. Oneextra GO list was added: Epigenomics_Roadmap_HM_ChIP-seq.gmt. For the Lemonite track, peptide abundance tables were generated in DIANN and exported for protein-level statistical inference using MSqRob2^15^. This split design for protein quantification underlies smaller discrepancies between the two perspectives.

### 6. Metabolome data processing

The result file from Compound Discoverer was imported into MetaX for data preprocessing and further analysis. Preprocessing included normalizing the data to obtain relative peak areas by Probabilistic Quotient Normalization, PQN. Quality control-based robust LOESS signal correction was used to correct Batch effects. Metabolites with a Coefficient of Variation larger than 30% on their Relative peak area in QC Samples were removed. An overall reference vector was obtained by analyzing the ion intensity distribution in each sample. Analyze the correction coefficient between the actual sample and the reference vector for actual sample correction. QC-RLSC is a data correction method in the field of metabolomics, the method is able to correct experiment sample signal by local polynomial regression fitting signal correction only based on the QC sample.

### 7. Histone data preprocessing in Progenesis QIP

Mass-spectrometry–based approaches allow untargeted and more comprehensive profiling of (combinatorial) hPTMs, and are particularly suited to investigate the metabolic read-out of the energy-information axis. Readers are encouraged to consult ^20,35,46–48^, for a more detailed rationale, technical considerations, and overall analytical framework underlying sample extraction, mass spectrometry data acquisition, and post-acquisition data analysis, which were adapted in the present study. Provez et al. ^35^ is a tutorial for the manual curation of detected histone features in a publicly accessible Progenesis QIP project, equally allowing to import additional searches to extend the current hPTM repertoire of this study *ad libidum*.

Briefly, raw data from all runs were imported and all runs were aligned in a single experiment in Progenesis QIP 4.2 (Nonlinear Dynamics, Waters) for feature detection. Next, the 20 MS/MS spectra closest to the elution apex were selected for each precursor ion and merged into a single ∗.mgf file to search in Mascot (Matrix Science). Mascot is a database search engine that uses a probabilistic scoring algorithm to define peptide-to-spectrum matches (PSM) and does not rely on target-decoy searched for FDR, as these are not applicable in combinatorial PTM search spaces. Two types of searches were performed on this file: 1) a quality search to verify the presence of non-propionylated standards (ß-gal), and to assess the extent of underpropionylation; and 2) an error tolerant search to identify the proteins present in the sample, and to determine the best set of 9 hPTMs for further analysis. The Mascot search parameters used for both searches are shown in Table 1. A smaller FASTA-database containing all identified proteins, including the acid extractome, was generated based on the results from the error tolerant search for further analysis. Next, the three MS/MS spectra closest to the elution apex were merged into a single *.mgf and exported to perform a biological search in Mascot (Table 1). The search results (∗.xml-format) were parsed to remove fixed propionylations and abbreviate the variable modifications and imported back into Progenesis QIP 4.2 to annotate the features from which they originated. Features that were annotated as histone peptidoforms were manually validated by an expert to resolve isobaric near-coelution. These are tagged as “edited”. Consequently, runs were normalized against all histone peptides in order to assure a constant histone protein abundance, since we aim to quantify changes in hPTMs and not in the expression of histones themselves.

Outliers were removed based on normalization factor (greater than 3 and less than 0,3) and PCA clustering of all histone peptidoforms. Finally, deconvoluted peptide ion data (*.csv format) from all histones were exported from Progenesis QIP 4.2 for statistical inference to PTM level in a histone adaptation of MSqRob2PTM^15,35^. A detailed description is given below. Further analysis and visualisation of results, including heatmaps, was performed using custom R scripts.

### 8. Proteome data analysis

Analysis of proteomics DIA data was performed using DIA-NN (version 18.1) 275. Raw files were imported into DIA-NN, and preprocessing steps included noise reduction and peak picking. A spectral library was generated from a FASTA file containing human protein sequences retrieved from Uniprot (accessed on Dec 6th, 2023), supplemented with contaminants. Peptide identification was performed using a combination of library matching and deep neural network prediction using double-pass mode as neural network classifier. Quantification was performed using the robust LC high accuracy setting. Mass accuracy was set to 20 ppm and MS1 accuracy was set to 10 ppm. Finally, peptide abundance tables were generated and exported to (i) Omics Playground (Bigomics) for the proteome and multiomics reports and (ii) to MSqRob2 (https://github.com/statOmics/msqrob2) for more advance protein-level statistical inference for downstream Lemonite analysis and to build the biomolecular network.

Summarized single PTM values and corresponding summarized protein abundances for each sample were correlated by Pearson correlation using a customized R script. For each PTM, the resulting correlation coefficients were ranked to generate a rank plot, highlighting proteins with the strongest positive and negative correlations.

### 9. Msqrob2PTM for Histones

The histone workflow applied is unique and its conceptual framework is described in more detail elsewhere ^16^. Briefly, hPTMs are fundamental to epigenetic regulation, yet their quantitative analysis from MS data remains challenging. The conventional method of calculating relative abundance (RA) to infer hPTM abundance has been instrumental, yet suffers from significant limitations, including a lack of robustness against missing data, biases from differential ionization efficiencies, and normalization artifacts in case of limited peptidoform coverage. Here, we therefore implemented a novel application of the msqrob2PTM statistical framework for the differential usage analysis of hPTMs, adding in histone-tailored functionalities and using it to address the shortcomings of RA (**Supplementary Figure 7**). Our workflow begins by establishing unified hPTM definition across highly conserved histone variants using multiple sequence alignment (MSA) to resolve positional ambiguity. We then introduce two complementary analysis strategies: a variant-corrected workflow that normalizes peptidoform abundances against their variant group to investigate variant-specific hPTM usage, and a variant-agnostic workflow that normalizes against total histone abundance returning a combined measure of variant and hPTM effects. By extending the analysis to include differential variant usage, we additionally provide a multi-layered view of the histone code, aiding in the interpretation of results. This flexible framework presents a statistically sound and robust alternative to RA, advancing our ability to analyze MS-based histone data. The html report contains all code for reproduction and customizations.

### 10. MSA and hPTM Definition

Individual .fasta files were downloaded from UniProt ^49^ (2026-02-18 17.52h) for every histone family using queries containing organism_id:9606, reviewed:true, and family:"histone x family" where x was one of H1/H5, H2A, H2B, H3, or H4. These sequences were added into the corresponding histone family reference sequence alignment retrieved from HistoneDB 2.0 ^50^ using the --add functionality of MAFFT (v 7.525). Histone peptidoforms were then mapped to all potential variants of origin, the positions of hPTMs were then translated for each matching variant to the aligned position within the MSA and finally mapped onto the corresponding aligned canonical histone defined in HistoneDB 2.0, such that insertions in respect to the canonical sequence received a positional suffix of “.X” where X is the index of the residue in the insertion and residues before the canonical initiator methionine received a negative position counting down from the initiator methionine. Finally, modifications that were quantified through the same set of peptidoforms were grouped together into hPTM groups, which could then be inferred using the msqrob2PTM workflow (**Supplementary Figure 7**).

An extensive interactive HTML report is available at http://www.lemonite.ugent.be/AML_Results/.. with detailed explanations on each step of the analysis. The code for the entire workflow can be downloaded from the report (upper right corner).

An adapted workflow based on msqrob2PTM^15^ was used. Peptidoform abundances were normalized for technical variation using variance stabilizing normalization ^51^ and peptidoforms were filtered to remove contaminants and peptidoforms quantified in less than half of the samples or 3 samples in total, whichever was most strict. Histone peptidoforms were then usage normalized by removing either the summarized abundance of all histone peptidoforms (variant-agnostic, the alternative to all peptidoforms of the corresponding variant group in the variant-corrected workflow), in turn calculated through robust summarization ^15^. Variant usage was then calculated through robust summarization of variant-agnostic peptidoform usage values corresponding to the given variant group, and hPTM (group) usage was calculated through robust summarization of variant-agnostic peptidoform usage values of peptidoforms containing the given hPTM group. Note, that if peptidoforms carried multiple PTMs, they would be used in multiple summarizations, once for each unique hPTM they carried. In turn, this implies that if two hPTMs are only detected on single peptidoform, they will have identical quantification outcomes.

The differential usage analysis was now carried out using the msqrob2 framework, which functions by modeling the usage normalized peptidoform/hPTM/variant abundances according to the different conditions specified by the experimental design. In this case, the 18 experimental groups were the 18 AML cell lines, which were modeled following the formula ∼ 0 + group. By testing the relevant contrasts, the difference can be evaluated. Peptidoforms, hPTMs, and variants were deemed significantly differential at a Benjamini–Hochberg adjusted P-value < 0.05.

### 11. Proteomics data processing

Raw protein abundance values were loaded and mapped to gene symbols using the UniProt ID Mapping portal (https://www.uniprot.org/id-mapping/). Non-unique gene symbols were discarded. Sample outliers identified in a prior dataset were removed before downstream analysis. Missing values were imputed with zero. Protein abundances were row-wise z-score normalized across samples. Normalized data were written to file as the basis for all subsequent analyses.

### 12. Metabolomics data Preprocessing

Two untargeted metabolomics datasets—acquired by HILIC (HI) and reversed-phase C18 chromatography—were processed in parallel. Only metabolites with level-1 or level-2 spectral confidence annotations were retained. Metabolite names were standardized to lowercase with special characters (spaces, hyphens, colons, parentheses, plus signs) replaced by underscores, and each metabolite was suffixed with _HI or _C18 to indicate dataset of origin. A log(x + 1) transformation was applied to each dataset independently. The two datasets were then concatenated by row-binding into a single metabolomics matrix, which was written to file together with a metabolite name–HMDB ID mapping table.

### 13. hPTM data Preprocessing

Histone PTM relative abundances were loaded from mass spectrometry-based histone proteomics data. Special characters in PTM identifiers (spaces, dashes, colons, and plus signs) were replaced with underscores. Sample identifiers were harmonized with the proteomics sample naming convention and sample sets were intersected to retain only common samples. Row-wise z-score scaling was applied.

### 14. Dimensionality reduction and quality control

All three omics layers—proteomics, histone post-translational modifications (hPTMs), and metabolomics—were subjected to quality-control visualization and dimensionality reduction (Preprocessing_and_TFA_clean.R). For each layer, per-feature z-score normalization (proteomics, hPTMs) or Pareto scaling (metabolomics, via IMIFA::pareto_scale()) was applied after initial preprocessing. Distribution boxplots were generated before and after normalization. Principal component analysis (PCA, stats::prcomp()), Uniform Manifold Approximation and Projection (UMAP, umap R package), and t-distributed Stochastic Neighbour Embedding (t-SNE, Rtsne) were performed to visualize sample structure. Dimensionality reduction quality was assessed quantitatively by the between-cluster to within-cluster sum-of-squares ratio (BCSS/WCSS), grouping samples by cell line identity.

### 15. Multi-omics network inference using Lemonite

Preprocessed proteomics, hPTM, and metabolomics matrices were formatted and provided as input to the LemonTree algorithm (v3.1.1), executed via the Lemonite Nextflow pipeline (http://www.lemonite.ugent.be). LemonTree identifies co-expression modules by iterative Gibbs sampling over a Bayesian network model linking co-expressed gene clusters to candidate regulators. One hundred independent Gibbs sampling runs were performed. Separate LemonTree runs were executed for: (i) the joint proteomics + hPTMs + metabolomics dataset (multiomics), (ii) metabolomics features alone, and (iii) hPTM features alone. For each run, LemonTree outputs a tight_clusters.txt file (gene-to-module assignments), an allreg.txt file (all regulator-to-module scores), a topreg.txt file (top 1% regulators), and a randomreg.txt file (scores from random regulatory associations).

#### Regulator selection

Regulators were selected from the allreg.txt output using a fold-over-random criterion (LemonTree_to_network.ipynb). For each regulator-module pair, the association score was compared against the maximum random score observed across all modules (derived from randomreg.txt). Regulators with a score ≥ 2× the global maximum random score (fold cutoff = 2) were retained. This was applied independently for hPTM regulators and metabolite regulators. Selected regulator-module scores were subsequently globally sum-normalized: each score was divided by the total sum of all selected scores and multiplied by 1000, making scores comparable across regulator types and rounding to the nearest integer.

#### Module coherence filtering

All LemonTree modules were evaluated for co-expression coherence (LemonTree_to_network.ipynb). For each module, a module eigengene was computed as the first principal component (PCA) of the module’s feature expression matrix across samples. Gene-module membership (kME) was defined as the absolute Pearson correlation of each gene’s expression profile with the module eigengene. Module coherence was defined as the mean absolute kME across all genes in the module. Modules with coherence < 0.5 or containing fewer than three genes were removed. The list of retained modules was saved to Networks/specific_modules.txt. Modules were ranked from highest to lowest coherence for downstream prioritization, and coherence scores were exported to Module_coherence_scores.txt.

#### Module pathway enrichment

For each retained module, a gene ranking was constructed by computing Pearson correlations between all proteins in the lemonite network and the module eigengene (calculated using WGCNA::moduleEigengenes()). GSEA was performed on this ranking using clusterProfiler (GO Biological Process, Molecular Function, Cellular Component, and all ontologies combined) and ReactomePA (Reactome), as well as KEGG and WikiPathways (Enrichment_on_modules_and_targets_full_dataset.R). A Benjamini–Hochberg FDR threshold of 0.05 was applied. The top 10 most significant up- and down-regulated pathways per module per database were collected and saved as summary CSV files. Module overview tables combining gene lists, hPTM and metabolite regulators, and pathway enrichment results were generated with Module_Overview_Generator_with_megaGO.ipynb, using the coherence-filtered module set. An interactive HTML summary report was generated from all output files using a custom Python script.

### 16. Correlation analysis – functional entanglement in full cellular context

#### Pairwise Pearson Correlation Analysis

All pairwise Pearson correlation coefficient (PCC) matrices were calculated across the three omics layers using preprocessed data (Generate_all_PCCs.R). Six inter- and intra-omics comparisons were computed: metabolites × proteins, metabolites × hPTMs, metabolites × metabolites, hPTMs × proteins, hPTMs × hPTMs, and proteins × proteins. Correlations were computed using stats::cor() with method = "pearson" and use = "pairwise.complete.obs". For visualization, features with the highest variance were prioritized in heatmaps rendered with pheatmap. Distribution plots of PCC values were generated as histograms. Summary statistics (mean, median, range) were reported for each matrix. All matrices were exported as CSV files.

#### Correlation network construction

Intra-omics co-correlation networks were constructed from the pairwise PCC matrices (Generate_correlation_networks.R). Edge lists were filtered by absolute PCC thresholds: |PCC| ≥ 0.7 for hPTM–hPTM networks, |PCC| ≥ 0.8 for all-metabolite networks, and |PCC| ≥ 0.7 for networks restricted to HMDB-annotated metabolites. Network node attributes included modification type (for hPTMs, extracted from the PTM identifier string) and metabolite superclass (from HMDB classification). Node and edge attribute tables were exported in tab-separated format compatible with Cytoscape.

#### Interactive network visualization

Correlation network node and edge attribute files were read and converted to interactive HTML visualizations using Cytoscape.js (visualize_correlation_networks_cytoscapejs.py, Python ≥ 3.8). Edge width was scaled proportionally to absolute PCC. Node colors were assigned by modification type (hPTMs) or metabolite superclass, using a fixed categorical color palette. One self-contained HTML file was produced per network and can be opened in any modern web browser without additional dependencies.

#### Gene Set Enrichment Analysis on Protein Correlation Profiles

For each hPTM, metabolite, or protein feature, a ranked gene list was constructed from its Pearson correlation profile across all detected proteins. Gene symbols were converted to Entrez IDs using clusterProfiler::bitr(). Gene Set Enrichment Analysis (GSEA) was performed with clusterProfiler (GO Biological Process, Molecular Function, Cellular Component, KEGG, WikiPathways) and ReactomePA (Reactome) (PCC_hPTM_protein_gsea.R, PCC_metabolite_protein_gsea.R, PCC_protein_protein_gsea.R). GSEA was parallelized across features using the future/future.apply framework (six workers). Random seed was set to 1234 for reproducibility. Results were saved per-feature in individual subdirectories together with pathway enrichment plots.

#### Module eigengene correlation

Module eigengenes were computed for independently inferred hPTM and metabolite co-expression modules using WGCNA::moduleEigengenes (correlation_hPTM_metabolite_modules.R). The first principal component of each module’s feature matrix was used as the eigengene. Pairwise Pearson correlations were then computed between all metabolite module eigengenes and all hPTM module eigengenes across samples. The resulting correlation matrix was visualized as a hierarchically clustered heatmap with pheatmap.

## DATA AVAILABILITY

All data can be interrogated through the http://www.lemonite.ugent.be/AML_Results. A video tutorial to help navigate the data is available at https://youtu.be/pSfDwsrRzm0. Upon registration to Omics Playground (Bigomics) two **Public Datasets** are freely accessible: *AML_18CellLines_June2025* (proteomics) and *AML_18CellLines_June2025_Multi-omics* (with full multi-omics functionalities).

RAW data and Supplementary Data will be made publicly available in the peer-reviewed manuscript. An overview is provided in this MindMap.

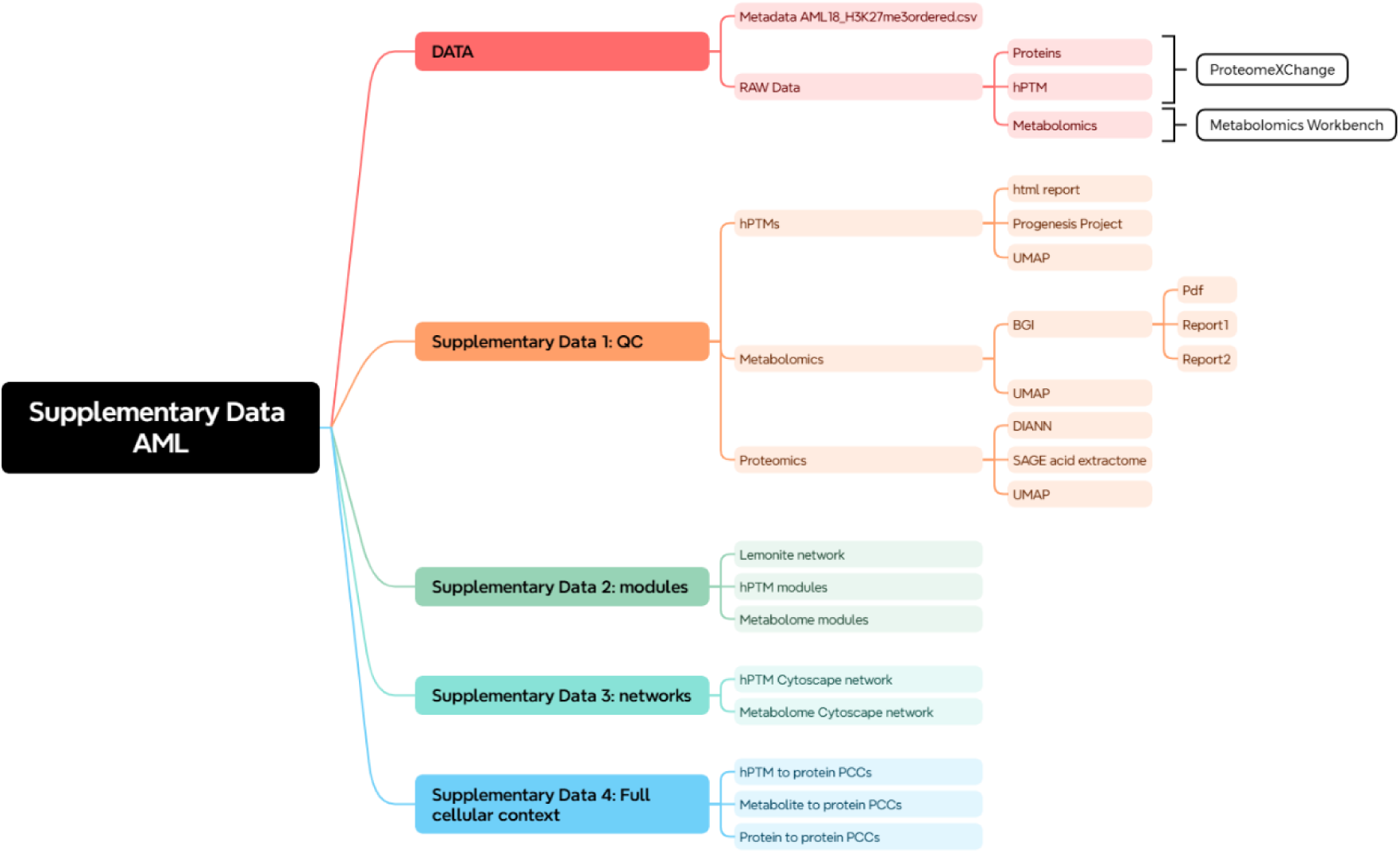

## DISCLAIMER

AZ and IK are employed by BigOmics Analytics at the time the work was conducted.

## ACKNOWLEDGEMENTS

This study has been supported by grants from The Research Foundation Flanders (FWO) awarded to L.C. (1SF2622N). The ZenoTOF 7600 instrument was funded through an “FWO medium-scale research infrastructure” grant (I008822N). R.A. acknowledges the Ghent University Special Research Fund (BOF), code 01D07621. The authors like to acknowledge ProGenTOmics for generating the proteomics and histone data and BGI Genomics for measuring and analysing the metabolomics.

## SUPPLEMENTARY FIGURES

**Supplementary Figure 1:**
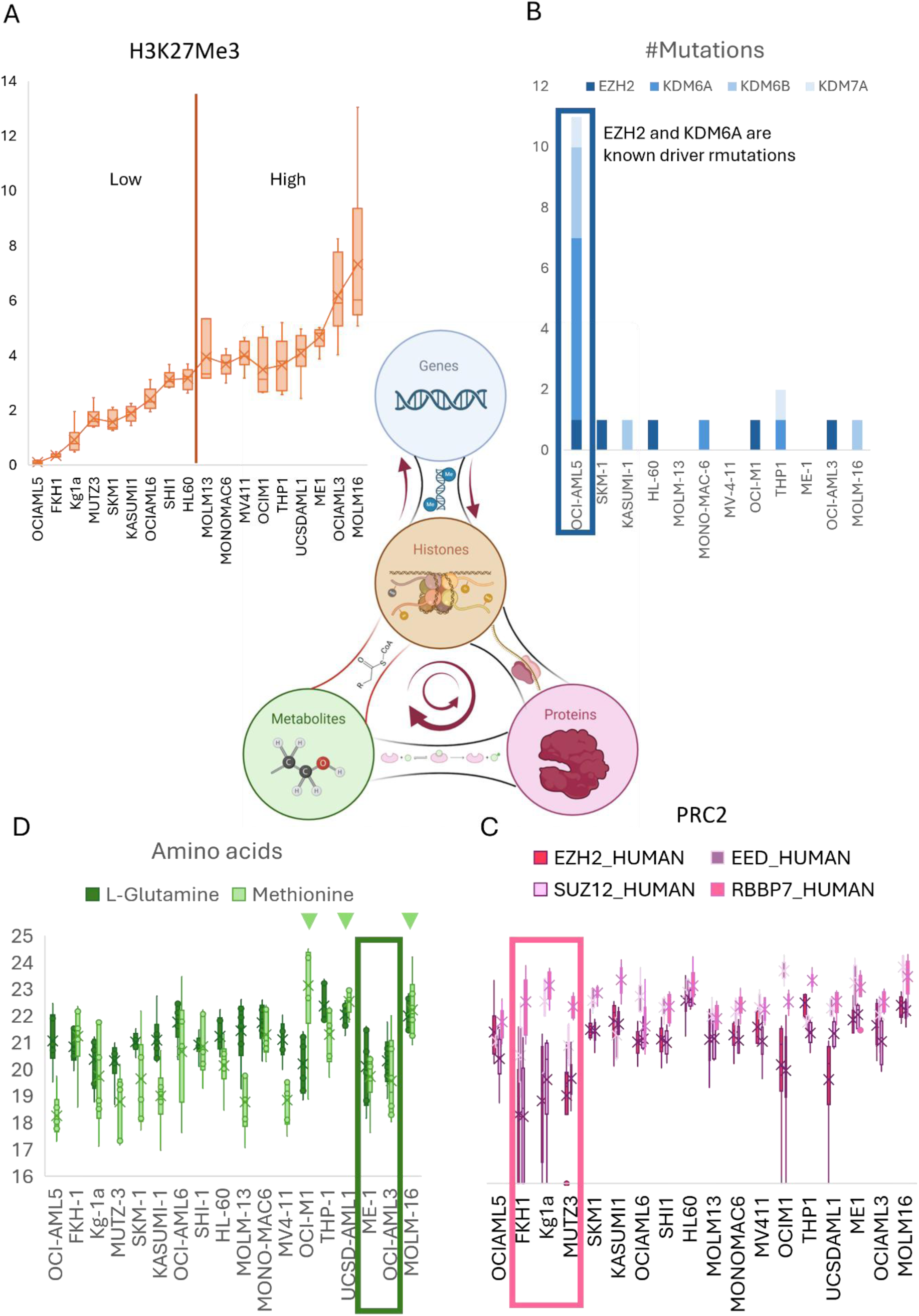
Illustrating the multi-omics picture from the targeted perspective of H3K27me3. The central cartoon depicts an extended version of the new biomolecular model from **Figure 1**. This equally provides a color-coded legend for the graphs. **A.** All cell lines are ordered according to their H3K27me3 levels (low to high) for all molecular fractions. Specific biomolecules from each fraction that potentially influence H3K27me3 are highlighted in **B-D** to illustrate the importance of capturing all these layers. **B.** Known mutations in writers and erasers of H3K27me3 taken from https://depmap.org/portal. The lowest H3K27me3 cell line has the most mutations. **C.** Expression of the four core units of the PRC2 complex which catalyzes the addition of this hPTM, as measured in the proteome fraction. For three out of four of the H3K27me3-lowest cell lines, the stoichiometry of the different units in the complex are disrupted. **D**. hPTMs are linked to metabolites in at least two distinct ways. First, metabolites act as co-factors that can activate and deactivate epigenetic readers and erasers. One example is glutamine, which is the precursor of a-ketoglutarate, an important co-factor of the JMJD demethylases. In fact, in tumors, glutamine deficiency has been directly linked to increased H3K27me3 and subsequent dedifferentiation, which, again, is especially relevant to AML^52^. This direct link could underly the high H3K27me3 levels in ME-1 and OCI-AML3 lines, yet those also have low methionine levels. Indeed, secondly, certain metabolites serve as substrates for modifications, such as most prominently SAM, which is the sole donor for methylation, and which is produced from the essential amino acid methionine (with the highest H3K27me3 cell lines showing the highest concentration). Interestingly, dietary methionine starvation has shown to delay AML progression both in vitro and in vivo ^53^. These are only a few of many factors conspiring on the final H3K27me3 abundance.

**Supplementary Figure 3.**
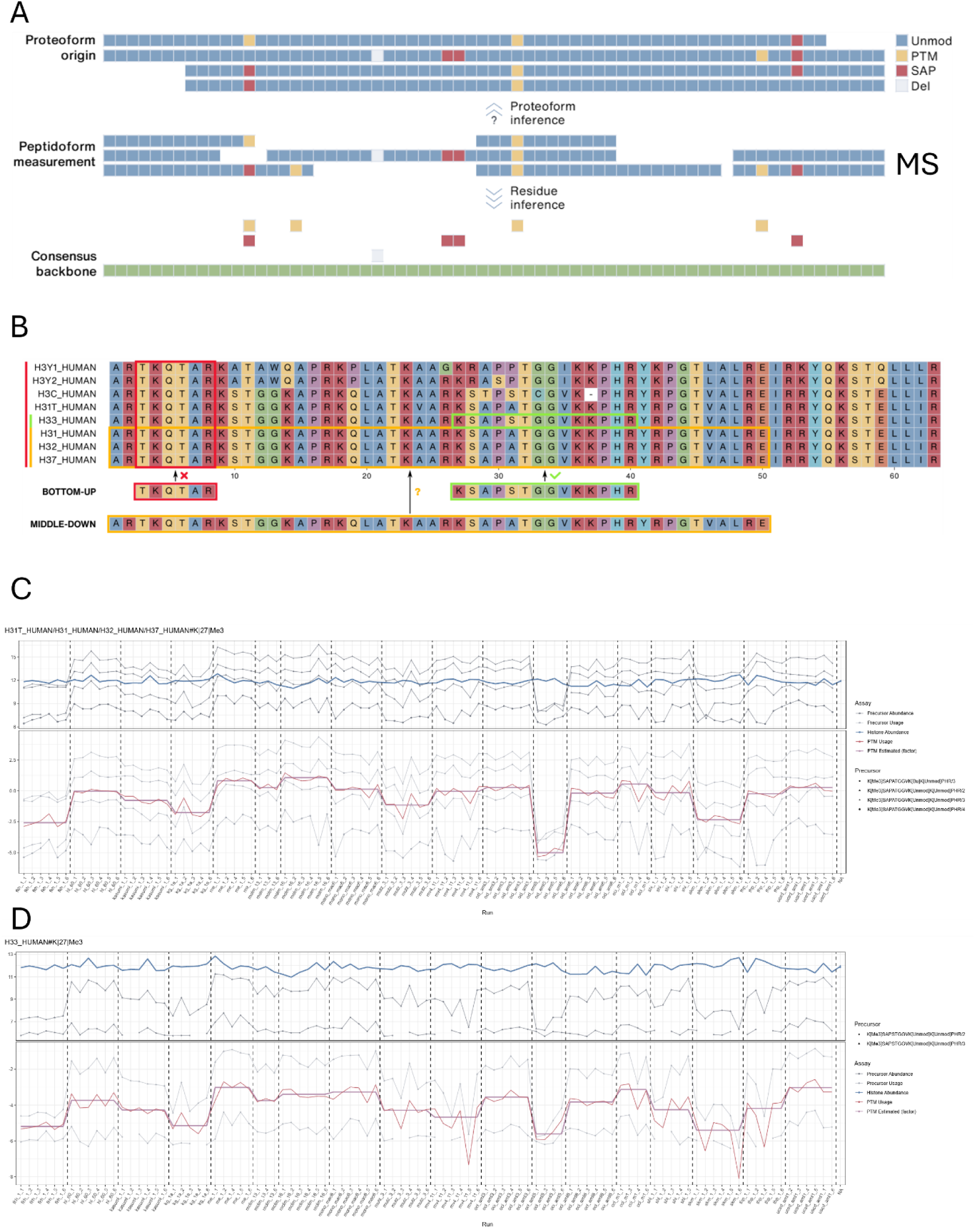
hPTM differential usage. **A.** The msqrob2PTM statistical framework was designed to robustly distinguish changes in PTM stoichiometry from changes in the abundance of the parent protein. To this end, it introduced the concept of differential PTM usage (DPU), which assesses PTM changes after correcting for parent protein abundance. The features measured by MS (middle) are the peptidoforms generated by trypsin digestion after and following by propionic anhydride derivatization. MSqRob uses these peptides to infer the protein abundance (upwards), while MSqRob2PTM infers hPTM abundance from the peptidoforms (downwards). Methodologically, preprocessed (e.g., filtering, global normalization, log₂-transformation) peptidoform abundances are first corrected by subtracting their corresponding parent protein abundance. PTM usage is then calculated through summarization, e.g., using the robust summarization procedure, of all peptidoforms containing a given PTM. **B.** To perform such DPU analysis, a relevant backbone is required to which usage can be defined: in msqrob2PTM, this backbone is the parent protein. This means that individual peptides must be inferred to their protein (group) of origin before abundances can be summarized and corrected. Despite the ambiguity inherent in protein inference, many proteotypic peptides can still be confidently inferred. This does not hold true for hPTM analysis; high sequence conservation makes it very difficult to unambiguously infer and quantify individual histone variants, i.e., the parent proteins of histone peptidoforms. This problem is most apparent for the well-studied H3 family: only few bottom-up (i.e., Arg-C specificity) histone peptides can be assigned to one H3 variant, and even some middle-down (i.e., Glu-C specificity) histone peptides are ambiguously inferred (yellow, not in the current data). Sequences of all human histone H3 variants (except CENPA_HUMAN, for simplicity) were added to the curated H3 multiple sequence alignment using MAFFT 7^54^ and the aligned sequences were colored based on the Clustal X color scheme, which categorizes amino acids by their physicochemical properties. Here, H3K_27_-R_40_ is the only stretch that can distinguish H31 and H33 after the applied bottom-up workflow. **C and D.** Differential usage plots can be downloaded from the report to manually inspect the quantitative data used to model the hPTM trends. H3.1K_27_-R_40_ (C) and H3.3K_27_-R_40_ (D) are depicted. The top box of each plot depicts the measured peptidoforms that contain the K27me3 hPTM and the blue line is the inferred H3 protein abundance. The lower box of each plot depicts the precursor usage normalized to the protein abundance and the thin red line depicts the inferred hPTM abundance. The bold purple line captures the modelled trend of the hPTM.

**Supplementary Figure 4.**
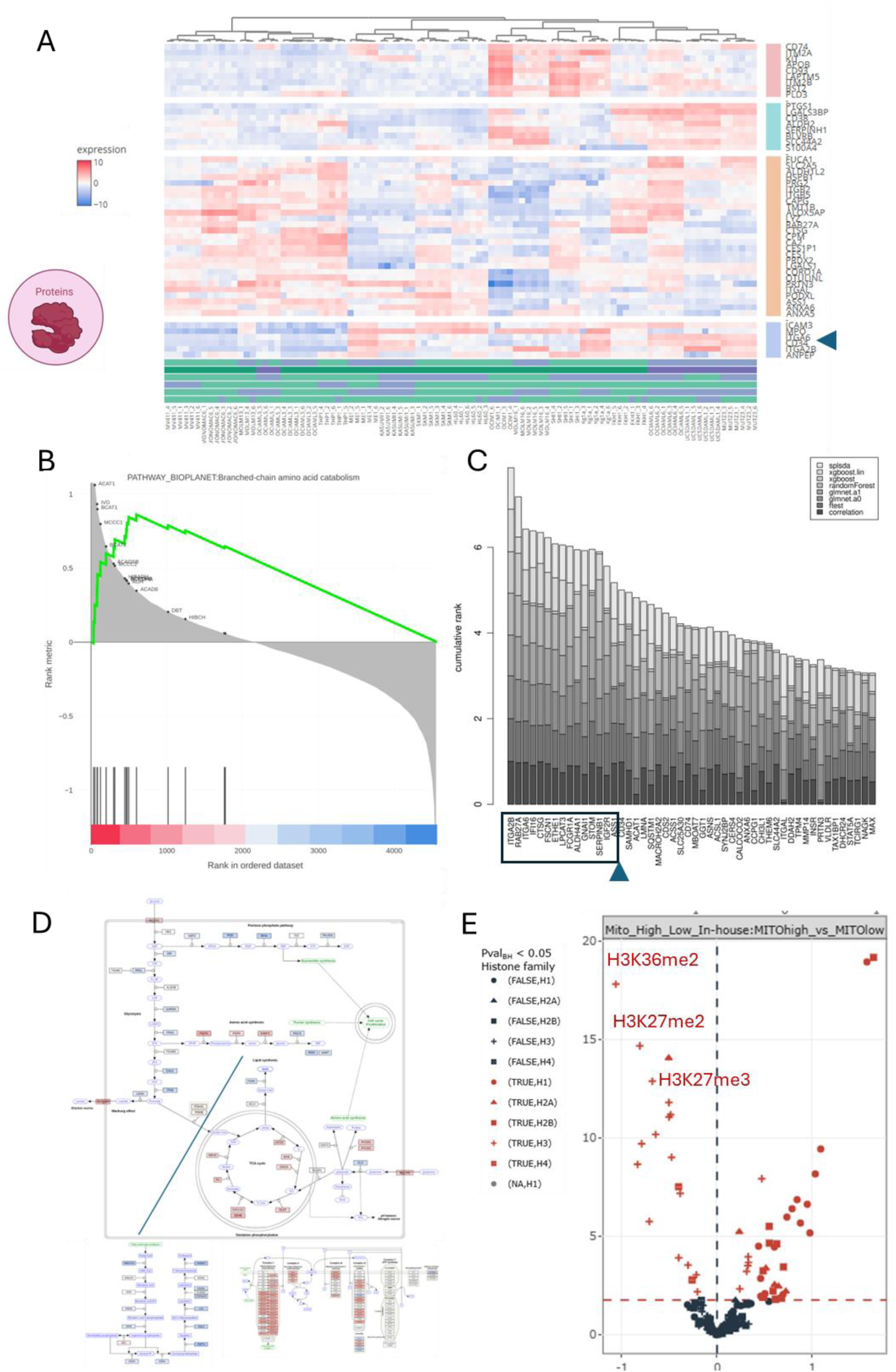
A The Clustered Heatmap with tree-like dendrogram shows the ‘distance’ between features and the approximate groups. This data-driven hierarchical clustering of the 18 cell lines shows that CD34 belongs to the 50 most resolving proteins in the measured proteome. **B.** GSEA analysis of the binary comparison between cell lines classified as C34+ vs CD34-, as seen by flow cytometry. The first hit is branched-chain amino acid catabolism. **C.** Machine learning-based biomarker discovery performed in Omics Playground (Bigomics). 15 proteins are considered better biomarkers than CD34 itself: : ITGA2B, RAB27A, ITGA6, IFI16, CTSG, FSCN1, ETHE1, LPCAT3, FCGR1A, ALDH4A1, GNAI1, STOM, SERPINB1, IGF2R and ASS1. Briefly, ITGA2B and ITGA6 are integrins, necessary for the interaction of myeloid cells with the bone marrow. Upregulated in CD34: (i) IFI16 (Gamma-interferon-inducible protein 16) may function as a transcriptional repressor and could have a role in the regulation of hematopoietic differentiation through activation of unknown target genes; (ii) ALDHA41, irreversibly converts delta-1-pyrroline-5-carboxylate (P5C), derived either from proline or ornithine, to glutamate. This is a necessary step in the pathway interconnecting the urea and tricarboxylic acid (TCA) cycles; (iii) ETHE1 (Persulfide dioxygenase ETHE1, mitochondrial) consumes molecular oxygen to catalyze the oxidation of the persulfide, once it has been transferred to a thiophilic acceptor, such as glutathione (R-SSH); (iv) LPCAT3 is a Lysophospholipid O-acyltransferase (LPLAT) that catalyzes the reacylation step of the phospholipid remodeling process also known as the Lands cycle. It catalyzes transfer of the fatty acyl chain from fatty acyl-CoA to 1-acyl lysophospholipid to form various classes of phospholipids. ASS1 (Argininosuccinate synthase) is downregulated in CD34 and is one of the enzymes of the urea cycle, the metabolic pathway transforming neurotoxic amonia produced by protein catabolism into inocuous urea. It catalyzes the formation of arginosuccinate from aspartate, citrulline and ATP and together with ASL it is responsible for the biosynthesis of arginine in most body tissues. **D.** GSEA analysis on the MitoHigh/MitoLow contrast shows very strong enrichment of Complex I, III and IV components of the respiratory chain. Strikingly, cholesterol metabolism is strongly inversely correlated in these cells. Mitochondrial function and cholesterol metabolism are usually tightly interdependent, and disruptions in one often disrupts the other. Little research supports a general negative correlation in which increased mitochondrial activity would systematically suppress cholesterol metabolism, or vice-versa. **E.** Volcano plot of the differential usage of hPTMs between Mito High (left) and Mito Low (right).

**Supplementary Figure 5.**
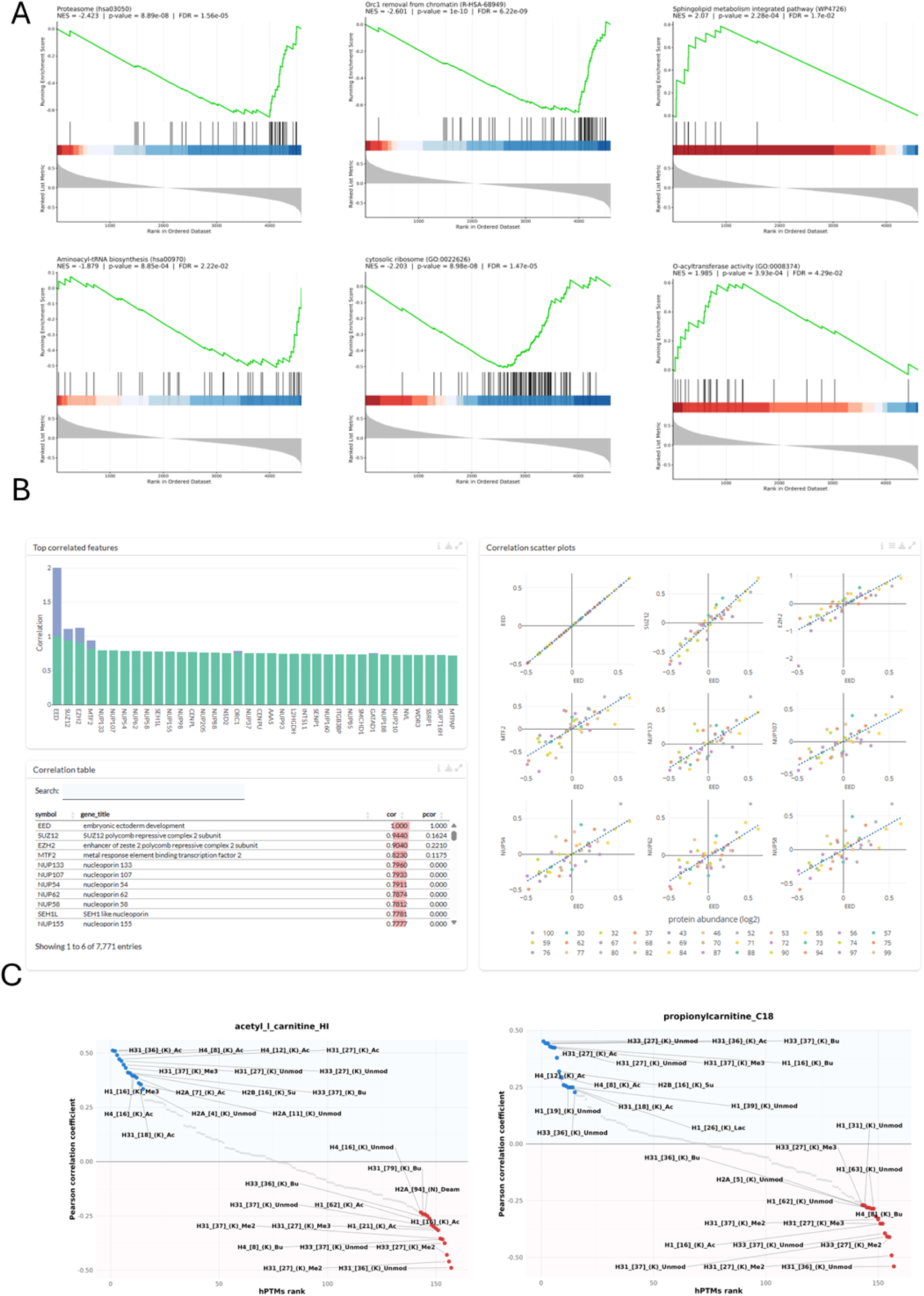
Targeting inter-omics edges. **A.** Positive and negative GO terms from the functional entanglement analysis to CD34. **B.** The proteome data downloaded from Kramer et al. ^37^ was loaded into Omics Playground (Bigomics) and a snippet from the correlation analysis is shown. From the perspective of EED, its PRC2 partners and the nuclear pore members also rank highest out of 7771 entries, confirming the correlations described in this manuscript. **C.** Ranked correlation plots of two more acyl-carnitines, showing the same functional entanglements as shown in **Figure 5H**.

## REFERENCES

1. Philip Ball. How Life Works. (The University of Chicago Press, 2023).

2. Noble, D. It’s time to admit that genes are not the blueprint for life. Nature 626, 254–255 (2024).

3. Klein, B., Hoel, E., Swain, A., Griebenow, R. & Levin, M. Evolution and emergence: higher order information structure in protein interactomes across the tree of life. Integrative Biology 13, 283–294 (2021).

4. van Mierlo, G. et al. Integrative Proteomic Profiling Reveals PRC2-Dependent Epigenetic Crosstalk Maintains Ground-State Pluripotency. Cell Stem Cell 24, 123–137.e8 (2019).

5. Zijlmans, D. W. et al. Integrated multi-omics reveal polycomb repressive complex 2 restricts human trophoblast induction. Nature Cell Biology 2022 24:6 24, 858–871 (2022).

6. Kumar, B. et al. Polycomb repressive complex 2 shields naïve human pluripotent cells from trophectoderm differentiation. Nat. Cell Biol. 24, 845–857 (2022).

7. Loh, J.-J. & Ma, S. Hallmarks of cancer stemness. Cell Stem Cell 31, 617–639 (2024).

8. Jayavelu, A. K. et al. The proteogenomic subtypes of acute myeloid leukemia. Cancer Cell 40, 301–317.e12 (2022).

9. Issa, G. C. et al. The menin inhibitor revumenib in KMT2A-rearranged or NPM1-mutant leukaemia. Nature 615, 920–924 (2023).

10. Kim, C. Y. et al. HumanNet v3: an improved database of human gene networks for disease research. Nucleic Acids Res. 50, D632–D639 (2022).

11. Brunk, C. F. & Martin, W. F. Archaeal Histone Contributions to the Origin of Eukaryotes. Trends in Microbiology Preprint at 10.1016/j.tim.2019.04.002 (2019).

12. Mesbahi, Y., Trahair, T. N., Lock, R. B. & Connerty, P. Exploring the Metabolic Landscape of AML: From Haematopoietic Stem Cells to Myeloblasts and Leukaemic Stem Cells. Front. Oncol. 12, (2022).

13. Katada, S., Imhof, A. & Sassone-Corsi, P. Connecting threads: Epigenetics and metabolism. Cell Preprint at 10.1016/j.cell.2012.01.001 (2012).

14. Vandemoortele, B. et al. Lemonite: identification of regulatory metabolites through data-driven, interpretable integration of transcriptomics and metabolomics data. Preprint at 10.64898/2026.03.27.714373 (2026).

15. Demeulemeester, N. et al. msqrob2PTM: Differential Abundance and Differential Usage Analysis of MS-Based Proteomics Data at the Posttranslational Modification and Peptidoform Level. Molecular & Cellular Proteomics 23, 100708 (2024).

16. Almey, R. Silencing the noise… Addressing common artifacts in histone proteomics. (Ghent University, 2025).

17. Ishikawa, K., Makanae, K., Iwasaki, S., Ingolia, N. T. & Moriya, H. Post-Translational Dosage Compensation Buffers Genetic Perturbations to Stoichiometry of Protein Complexes. PLoS Genet. 13, e1006554 (2017).

18. Meert, P., Govaert, E., Scheerlinck, E., Dhaenens, M. & Deforce, D. Pitfalls in histone propionylation during bottom-up mass spectrometry analysis. Proteomics 15, 2966–2971 (2015).

19. Meert, P. et al. Tackling aspecific side reactions during histone propionylation: The promise of reversing overpropionylation. Proteomics 16, 1970–4 (2016).

20. Willems, S. et al. Flagging False Positives Following Untargeted LC-MS Characterization of Histone Post-Translational Modification Combinations. J. Proteome Res. 16, 655–664 (2017).

21. Govaert, E. et al. Extracting histones for the specific purpose of label-free MS. Proteomics 16, 2937–2944 (2016).

22. Ball, P. Should biology put complexity first? Cell Syst. 16, 101197 (2025).

23. Hong, Y.-G. et al. The RNA m6A Reader YTHDF1 Is Required for Acute Myeloid Leukemia Progression. Cancer Res. 83, 845–860 (2023).

24. Huang, F. et al. m6A/IGF2BP3-driven serine biosynthesis fuels AML stemness and metabolic vulnerability. Nat. Commun. 16, 4214 (2025).

25. Tsitsiridis, G., et al. CORUM: the comprehensive resource of mammalian protein complexes–2022. Nucleic Acids Res. 51, D539–D545 (2023).

26. Steinkamp, R. et al. CORUM in 2024: protein complexes as drug targets. Nucleic Acids Res. 53, D651–D657 (2025).

27. Schuurhuis, G. J., et al. The Prognostic Value of CD34 Expression In Acute Myeloid Leukemia. A Mystery Solved. Blood 116, 2725–2725 (2010).

28. Hughes, M. R. et al. A sticky wicket: Defining molecular functions for CD34 in hematopoietic cells. Exp. Hematol. 86, 1–14 (2020).

29. Juluri, K. R., Siu, C. & Cassaday, R. D. Asparaginase in the Treatment of Acute Lymphoblastic Leukemia in Adults: Current Evidence and Place in Therapy. Blood Lymphat. Cancer Volume 12, 55–79 (2022).

30. Bartram, J. et al. Activation of a branched-chain amino acid rheostat restores replication-dependent hematopoietic stem cell fitness. Cell Stem Cell 10.1016/j.stem.2025.12.018 (2026) doi:10.1016/j.stem.2025.12.018.

31. Lesnik, C. et al. Enhanced branched-chain amino acid metabolism improves age-related reproduction in C. elegans. Nat. Metab. 6, 724–740 (2024).

32. Bonardi, F. et al. A Proteomics and Transcriptomics Approach to Identify Leukemic Stem Cell (LSC) Markers. Molecular & Cellular Proteomics 12, 626–637 (2013).

33. Tcheng, M. et al. Very long chain fatty acid metabolism is required in acute myeloid leukemia. Blood 137, 3518–3532 (2021).

34. De Clerck, L. et al. Untargeted histone profiling during naive conversion uncovers conserved modification markers between mouse and human. Sci. Rep. 9, 17240 (2019).

35. Provez, L. et al. An interactive mass spectrometry atlas of histone posttranslational modifications in T-cell acute leukemia. Scientific Data 2022 9:1 9, 1–15 (2022).

36. Gozalo, A. et al. Core Components of the Nuclear Pore Bind Distinct States of Chromatin and Contribute to Polycomb Repression. Mol. Cell 77, 67–81.e7 (2020).

37. Kramer, M. H. et al. Proteomic and phosphoproteomic landscapes of acute myeloid leukemia. Blood 140, 1533–1548 (2022).

38. Zijlmans, D. W. et al. PRC1 and PRC2 proximal interactome in mouse embryonic stem cells. Cell Rep. 44, 115362 (2025).

39. Chen, D. et al. Identification of Macrodomain Proteins as Novel O-Acetyl-ADP-ribose Deacetylases. Journal of Biological Chemistry 286, 13261–13271 (2011).

40. Dai, Z., Ramesh, V. & Locasale, J. W. The evolving metabolic landscape of chromatin biology and epigenetics. Nat. Rev. Genet. 21, 737–753 (2020).

41. Swanton, C. et al. Embracing cancer complexity: Hallmarks of systemic disease. Cell 187, 1589–1616 (2024).

42. Verhelst, S. & Corveleyn, L. Direct acid extraction v1. Preprint at 10.17504/protocols.io.bp2l6x6odlqe/v1 (2024).

43. Verhelst, S. One-dimensional SDS-PAGE (9-18% TGX gel) v1. Preprint at 10.17504/protocols.io.5jyl8pz27g2w/v1 (2024).

44. Corveleyn, L. & Verhelst, S. Propionylation and tryptic digestion v1. Preprint at 10.17504/protocols.io.5jyl8pz36g2w/v1 (2024).

45. Demichev, V., Messner, C. B., Vernardis, S. I., Lilley, K. S. & Ralser, M. DIA-NN: neural networks and interference correction enable deep proteome coverage in high throughput. Nat. Methods 10.1038/s41592-019-0638-x(2020) doi:10.1038/s41592-019-0638-x.

46. Daled, S. et al. Histone sample preparation for bottom-up mass spectrometry: a roadmap to informed decisions. Proteomes.

47. De Clerck, L. et al. An experimental design to extract more information from MS-based histone studies. *Mol*. Omics 17, (2021).

48. Verhelst, S. et al. A large scale mass spectrometry-based histone screening for assessing epigenetic developmental toxicity. Sci. Rep. 12, (2022).

49. Ahmad, S. et al. The UniProt website API: facilitating programmatic access to protein knowledge. Nucleic Acids Res. 53, W547–W553 (2025).

50. Draizen, E. J. et al. HistoneDB 2.0: a histone database with variants—an integrated resource to explore histones and their variants. Database 2016, baw014 (2016).

51. Huber, W., von Heydebreck, A., Sültmann, H., Poustka, A. & Vingron, M. Variance stabilization applied to microarray data calibration and to the quantification of differential expression. Bioinformatics 18, S96–S104 (2002).

52. Pan, M. et al. Regional glutamine deficiency in tumours promotes dedifferentiation through inhibition of histone demethylation. Nat. Cell Biol. 18, 1090–1101 (2016).

53. Cunningham, A. et al. Dietary methionine starvation impairs acute myeloid leukemia progression. Blood 140, 2037–2052 (2022).

54. Katoh, K. & Standley, D. M. MAFFT Multiple Sequence Alignment Software Version 7: Improvements in Performance and Usability. Mol. Biol. Evol. 30, 772–780 (2013).

